# MoDLE: High-performance stochastic modeling of DNA loop extrusion interactions

**DOI:** 10.1101/2022.04.13.488157

**Authors:** Roberto Rossini, Vipin Kumar, Anthony Mathelier, Torbjørn Rognes, Jonas Paulsen

## Abstract

DNA loop extrusion emerges as a key process establishing genome structure and function. We introduce MoDLE, a computational tool for fast, stochastic modeling of molecular contacts from DNA loop extrusion capable of simulating realistic contact patterns genome wide in a few minutes. MoDLE accurately simulates contact maps in concordance with existing molecular dynamics approaches and with Micro-C data, and does so orders of magnitude faster than existing approaches. MoDLE runs efficiently on machines ranging from laptops to high performance computing clusters, and opens up for exploratory and predictive modeling of 3D genome structure in a wide range of settings.

## Background

DNA loop-extrusion, in which DNA is progressively reeled into transient loops, emerges as a key process in genome structure and function. The growing list of cellular processes where loop-extrusion plays a critical role now includes transcriptional regulation [1,2], DNA repair [3], VDJ-recombination [4], and cell division [5]. Recent single-molecule imaging experiments have provided direct observations of loop extrusion in vitro [6,7].

High-throughput chromosome conformation capture sequencing, including Hi-C [8] and Micro-C [9,10], have advanced our abilities to map three-dimensional (3D) genome organization through quantification of spatially proximal genome regions. The resulting data is usually rendered as a matrix of intrachromosomal and interchromosomal contact frequencies. These data increasingly deepen our understanding of 3D genome organization, and show DNA loop extrusion as a key process shaping genome structure [11–13]. In fact, topologically associating domains (TADs), which show up as sub-megabase-sized domains covering most of higher eukaryote genomes, are formed by loop extrusion [12]. TADs are relevant units of gene expression regulation, and are associated with disease when disrupted [14].

DNA loop extrusion is carried out by ring-shaped proteins (including cohesin and condensin) belonging to the Structural Maintenance of Chromosomes (SMC) family. These proteins are often referred to as loop extrusion factors (LEFs) [15]. The exact mechanism of how loop extrusion takes place in interphase is not fully understood. There is, however, convincing evidence that SMCs bind DNA to perform ATP-dependent loop extrusion in a symmetric or asymmetric fashion. Recent evidence suggests that cohesin can extrude DNA with a “swing-and-clamp” mechanism [16], and in a nontopological configuration where DNA is not encircled by the cohesin ring [17,18]. A loop starts extruding when a LEF binds to a genomic region, and continues processively until it is stalled by a DNA-bound CCCTC binding factor (CTCF) oriented with its N-terminus pointing towards the extruding cohesin complex. A pair of CTCFs arranged in a convergent orientation can thus stall loop growth on both sides creating semi-stable loops visible in Hi-C as a characteristic “dot” at TAD corners [19]. Similarly, when extruding loops are stalled only on one side, a “stripe” can be observed along one or both TAD borders [20]. The protein WAPL transiently releases cohesin from chromatin, terminating the loop extrusion process [21,22]. The resulting loop-extrusion patterns have been found in a range of Hi-C datasets so far, emphasizing the evolutionary conserved role of loop extrusion in shaping 3D genome organization [19,23].

Disrupting any of the key proteins involved in DNA loop extrusion has a dramatic effect on genome 3D structure. WAPL depletion causes an increase in loop stability, with an accumulation of axial elements and weakening of compartments [21,24]. Depletion of cohesin causes a large fraction of TADs and loops to disappear [24–26]. Similarly, depletion of CTCF induces loss of loops and TADs [24,27].

Modeling and simulation of DNA-DNA contact patterns is a powerful approach for understanding underlying molecular mechanisms and for predicting the effect of DNA perturbations. Polymer simulations and molecular dynamics (MD) have been used for modeling of TADs to study their structure and dynamics [28–31]. Computational modeling and simulation of loop extrusion has proven useful for predicting the effects of perturbations to TAD borders, and to properly understand patterns seen in Hi-C data. Initial models [15,32] of loop extrusion used the Gillespie algorithm to characterize looping properties and chromatin compaction, and did not sample contact maps. Subsequent models used HOOMD particle simulation [33] to perform homopolymer simulations where modeled LEFs extrude the polymers and halt at boundaries with properties defined from CTCF motif instance orientation and ChIP-seq signal strength [11,25]. Recently, to efficiently simulate larger genome regions, a combination of one-dimensional (1D) simulations with 3D polymer modeling has been applied to sample multiple conformations combined into contact maps. LEF binding, release and stalling probabilities are then modeled explicitly [34–36]. These simulations are typically implemented using the OpenMM molecular simulation framework [37]. The simulations can be used to explore and rule out molecular mechanisms. For example, Banigan et. al assessed the level of DNA compaction that can be achieved by different loop extrusion mechanisms, and concluded that one-sided loop extrusion alone fails to achieve the level of compaction observed in large metazoan genomes [36]. Other approaches embed epigenetic data in combination with crosslinking proteins to model and study conformational variability across complex chromatin regions [38,39]. To the best of our knowledge, no standalone software for modeling and simulation of loop extrusion exists.

We introduce MoDLE (Modeling of DNA Loop Extrusion), a high-performance stochastic model of DNA loop extrusion capable of efficiently simulating contacts from loop extrusion genome wide. In contrast to MD simulation approaches, simulating loop extrusion contacts using MoDLE is a straightforward process only requiring two input files and execution through a command line interface (CLI). MoDLE can simulate a contact matrix with the molecular interactions generated by DNA loop extrusion on the entire human genome in a matter of minutes using less than 1 GB of RAM. Typical use cases include predicting Hi-C contact patterns from ChIP-seq (or similar) data, and predicting the effect of alterations, mutations, and structural variation to TAD borders. MoDLE opens up for rapid simulation and parameter exploration of DNA loop extrusion on genomes of any size, including large mammalian genomes.

## Results

### MoDLE: Modeling of DNA Loop Extrusion

MoDLE uses fast stochastic simulation to sample DNA-DNA contacts generated by loop extrusion. Binding and release of LEFs and barriers and the extrusion process is modeled as an iterative process (see Fig. 1). At the beginning of a simulation MoDLE goes through a burn-in phase where LEFs are progressively bound to DNA, without sampling molecular contacts. The burn-in phase runs until the average loop size has stabilized. Active LEFs are extruded through randomly sampled strides along the DNA in reverse and forward directions. Each epoch, LEFs are released with a probability based on the average LEF processivity and extrusion speed. LEFs that are released in the current epoch will rebind to randomly sampled DNA regions in the next epoch. Extrusion barriers (e.g. CTCF binding sites) are modeled using a two-state (bound and unbound) Markov process. Each extrusion barrier consists of a position, a blocking direction and the Markov process transition probabilities. The occupancy of each extrusion barrier can be specified individually through the score field in the input BED file. Alternatively, users can specify a uniform barrier occupancy that is applied to all extrusion barriers. MoDLE accepts a large number of optional parameters to specify the model’s behavior. For example, users can specify the number of LEFs to be instantiated for each Mbp of simulated DNA using the --lef-density parameter. LEF-barrier and LEF-LEF collisions are processed each simulation epoch. Collision information is used to update candidate strides to satisfy the constraints imposed by collision events, and to compute how extrusion in the next epoch should proceed.

**Fig 1:**
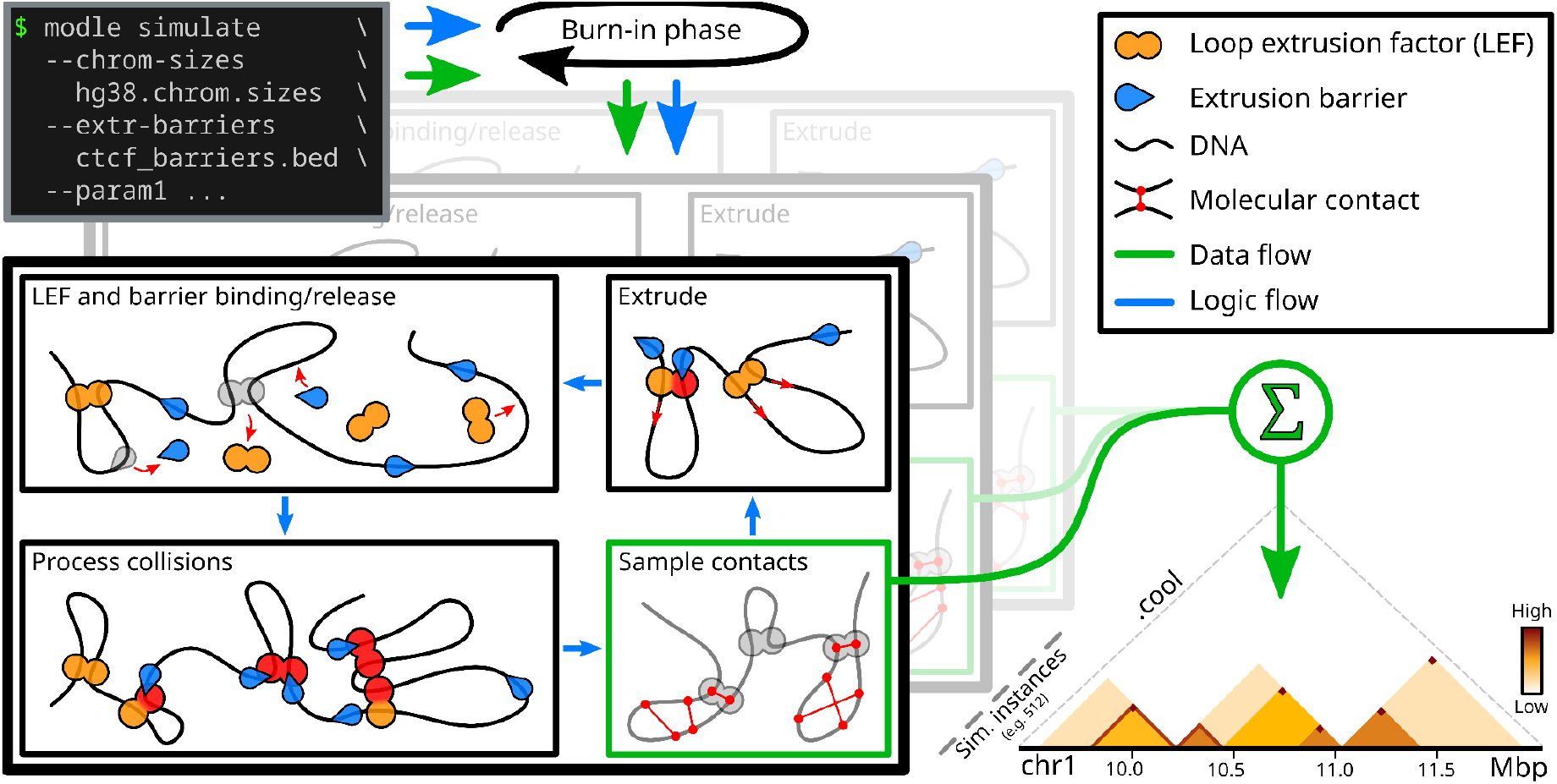
Schematic and simplified overview of MoDLE. Input files specify genome regions to be simulated (e.g. a chrom.sizes file) and their barrier positions (e.g. CTCF binding sites and orientation) in BED format. Optional parameters control the specifics of a simulation. Loop Extruding Factors (LEFs) bind to, extrude and release from the regions and interact with modeled barriers according to input parameters. Loop extrusion and intra-TAD contacts of a randomized subset of loops are recorded each epoch and aggregated into an output cooler file containing the final simulated contact frequencies. Simulation halts when a target number of epochs or a target number of loop extrusion contacts have been simulated.

During a simulation, sampled molecular contacts are accumulated into a specialized contact matrix data structure with low memory overhead. MoDLE execution continues until a target number of epochs or a target number of loop extrusion contacts are simulated. Finally, contacts generated by all simulation instances for a given chromosome are written to an output file in cooler format [40] (Fig. 1).

With default settings, MoDLE will run over 500 simulation instances for each chromosome simulated. Thus, simulation instances can run in parallel, making efficient use of the computational resources of modern multi-core CPUs. We designed MoDLE such that each simulation instance requires less than 10 MB of memory to simulate loop extrusion on large mammalian chromosomes, such as chromosome 1 from the human genome. To achieve high-performance, MoDLE stores most of its data in contiguous memory. Data is indexed such that extrusion barriers and extrusion units in a simulation instance can be efficiently traversed in 5’-3’ and 3’-5’ directions. This allows MoDLE to bind/release LEFs, process collisions, register contacts and extrude DNA in linear time-complexity.

More design and implementation details are available in Additional file 1 as well as MoDLE’s GitHub repository github.com/paulsengroup/modle.

### Comparison with Micro-C data and MD simulations

To assess MoDLE’s ability to reproduce contact data features known to be stemming from loop extrusion, we simulated genome-wide DNA-DNA contacts based on available CTCF and RAD21 ChIP-seq data in H1-hESC cells (see Methods). MoDLE is capable of simulating loop extrusion molecular contacts and intra-TAD contacts separately (see Section 9, Additional file 1 for details). A rendering of the resulting loop extrusion molecular contacts heatmaps show characteristic stripe and dot patterns at TAD borders (Fig. 2A). Simulated TAD contacts show enrichment of contacts within TADs, including a nested structure of the TADs (Fig. 2B). In combination, these patterns resemble well-characterized patterns observed in Micro-C and Hi-C data (Fig. 2C).

**Fig. 2:**
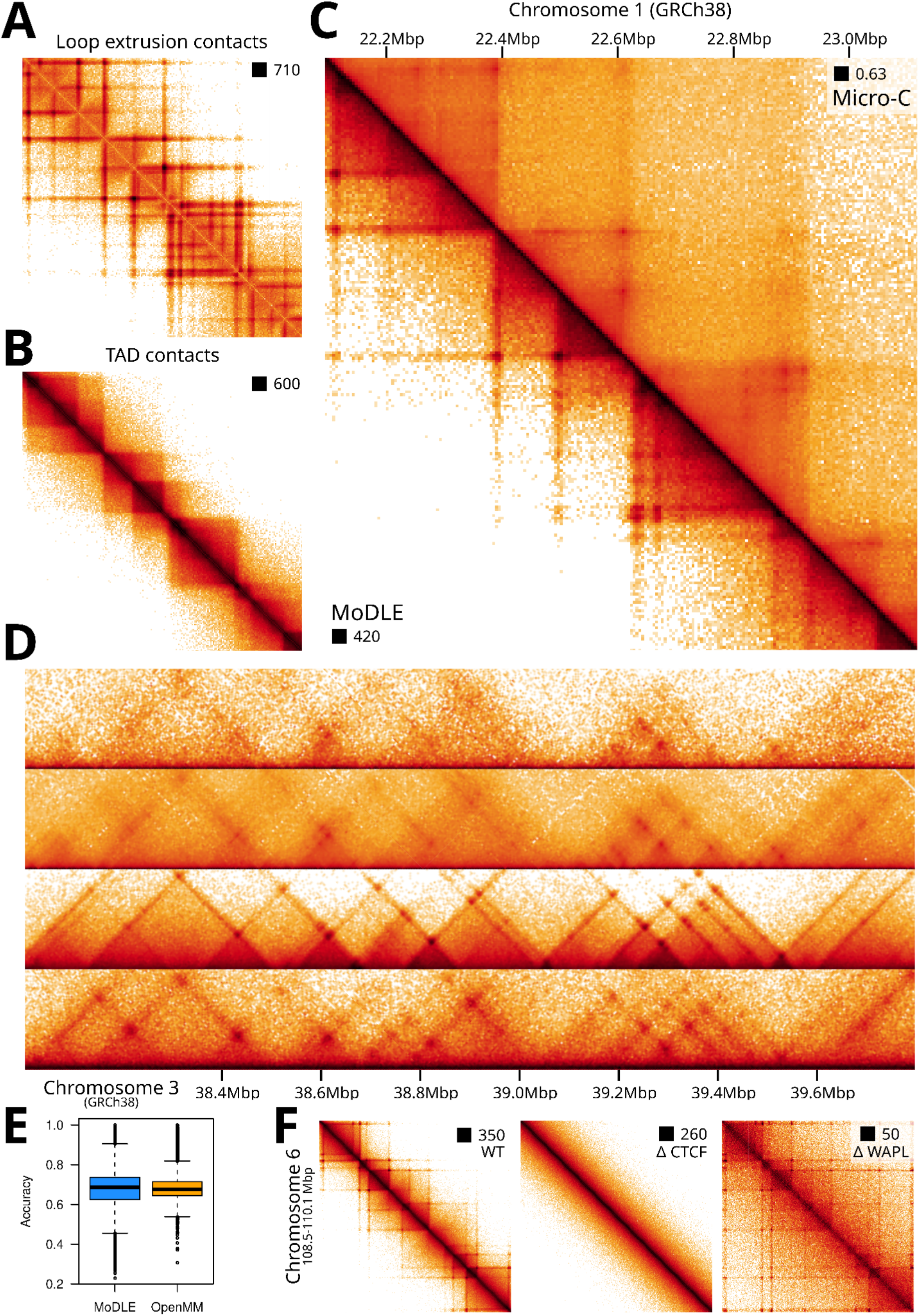
Comparison of MoDLE with OpenMM and Micro-C data. **A**: Simulated MoDLE contact frequencies solely mediated by LEFs. B: Intra-TAD contacts (only) generated with MoDLE. C: Lower triangle: Loop extrusion and intra-TAD contacts from MoDLE in the same region as for A and B. Upper triangle: Micro-C data from the same region. D: Side-by-side comparison of Micro-C data, MoDLE output and OpenMM output for a region on chromosome 3 in H1-hESC. E: Quantitative comparison of the accuracy (fraction of correctly classified pixels relative to all pixels) of MoDLE and OpenMM in reproducing stripe and dot pixel-patterns observed in modeled regions in H1-hESC cells (see Methods). F: *in silico* simulated molecular contacts mimicking CTCF and WAPL depletion. Left: Wildtype (WT) output of MoDLE in a region on chromosome 6 in H1-hESC. Middle: effect on MoDLE output when CTCF barriers weakly associate with their binding sites. Right: effect on MoDLE output when LEFs are less likely to be released from DNA, thus mimicking WAPL depletion.

Even though no stand-alone software exists for direct side-by-side comparison, we adapted available code based on OpenMM [36] to systematically compare the output with that of MoDLE (see Methods). We chose OpenMM for comparison as it is an efficient and widely used system for simulating loop extrusion [12,34–36,41].

Using the same input data, we simulated contacts in five different 10 Mbp regions on five different chromosomes. In general, MoDLE produces contact patterns similar to OpenMM (Fig. 2D and Supplementary Fig. 1, Additional file 2), and MoDLE output and OpenMM correlate strongly (Pearson p=0.93; see Supplementary Fig. 2, Additional file 2). By comparing contacts with corresponding Micro-C and Hi-C data (Fig. 2D), we see a median pixel accuracy (*i.e*. the ability to correctly classify pixels as a dot/stripe or not, relative to all pixels; see Methods) of 0.69 for MoDLE and 0.68 for OpenMM, signifying that MoDLE indeed simulates contacts observed in Micro-C similar to OpenMM (Fig. 2E). Note that contacts generated by OpenMM involve 3D polymer modeling and thus, unlike MoDLE, considers random polymer contacts. As a consequence, contacts not generated by loop-extrusion will be included in the OpenMM output. Therefore, long-range contacts (~2-3Mbp) are generally not as enriched in the MoDLE output as these contacts are mainly compartmental or dominated by random polymer interactions. This can be seen when employing a diagonal-by-diagonal correlation between MoDLE and OpenMM, which shows that the two methods correlate better at short range contacts than at long range contacts (see Supplementary Fig. 3). It implies that MoDLE does not by default recapitulate the relationship between the distance from the diagonal and the contact frequencies as seen in Hi-C or Micro-C data. However, when LEF processivity is increased, this trend is gradually approached (see Supplementary Fig. 4). Comparing the output of MoDLE and OpenMM in A and B compartments separately shows minimal difference of performance between compartments (Supplementary Fig. 5).

Altering MoDLE’s input parameters to *in silico* mimicking depletion of CTCF and WAPL, shows an expected loss of TAD insulation patterns [27] upon *in silico* depletion of CTCF, and more pronounced stripe and dot patterns [22] when mimicking WAPL depletion (Fig. 2F). Similarly, altering the parameters specifying LEF density, LEF processivity and LEF-LEF collisions shows relevant and predictable consequences in the data output (see Supplementary Fig. 6-10). We conclude that MoDLE is capable of simulating loop extrusion and TAD contact patterns similar to existing state-of-the-art molecular dynamics (OpenMM) approaches.

### Benchmarking of execution time and memory usage

MoDLE is designed for fast genome-wide simulation of loop extrusion contact patterns. A genome-wide run with default settings, simulating loop extrusion on the entire human genome using barriers from H1-hESC (38,815 CTCF barriers and 61,766 LEFs; see Methods) takes ~40 seconds on a compute server (server A; see Table 1) and ~5 minutes on a laptop (laptop A; see Table 1), generating over 370 million contacts. To systematically compare MoDLE execution time and memory usage with OpenMM, we generated synthetic input datasets with increasing genome size (1-500 Mbp) and number of CTCF barriers (4 barriers per Mbp of DNA simulated) (see Methods for details). The inputs were identical in MoDLE and OpenMM. Each measurement was repeated 10 times for MoDLE and 5 times for OpenMM. For MoDLE we run benchmarks using 1-52 CPU cores, while for OpenMM we tested the CPU (server C; see Table 1) and GPU (server D; see Table 1) implementations. We computed median elapsed wall clock time and peak memory usage for MoDLE and OpenMM. The resulting comparisons show that MoDLE simulations using 52 CPU cores complete within 0.7-71 seconds from the smallest to the largest genome region. OpenMM requires 2 hours and 35 minutes for the smallest genome region and over 41 hours for a genomic region of 250Mbp (Fig. 3A). Due to very long execution times, OpenMM runs above 250Mbp were not performed. For the compared genome regions, MoDLE is 4000-5000 times faster than OpenMM (Fig. 3A). OpenMM simulations without GPU acceleration were particularly slow and were only used to simulate genome regions below 5 Mbp, and required up to 35 hours 20 minutes of execution time (Fig. 3A). Thus, in practice running OpenMM requires access to GPUs, while MoDLE runs efficiently using CPUs.

**Fig. 3:**
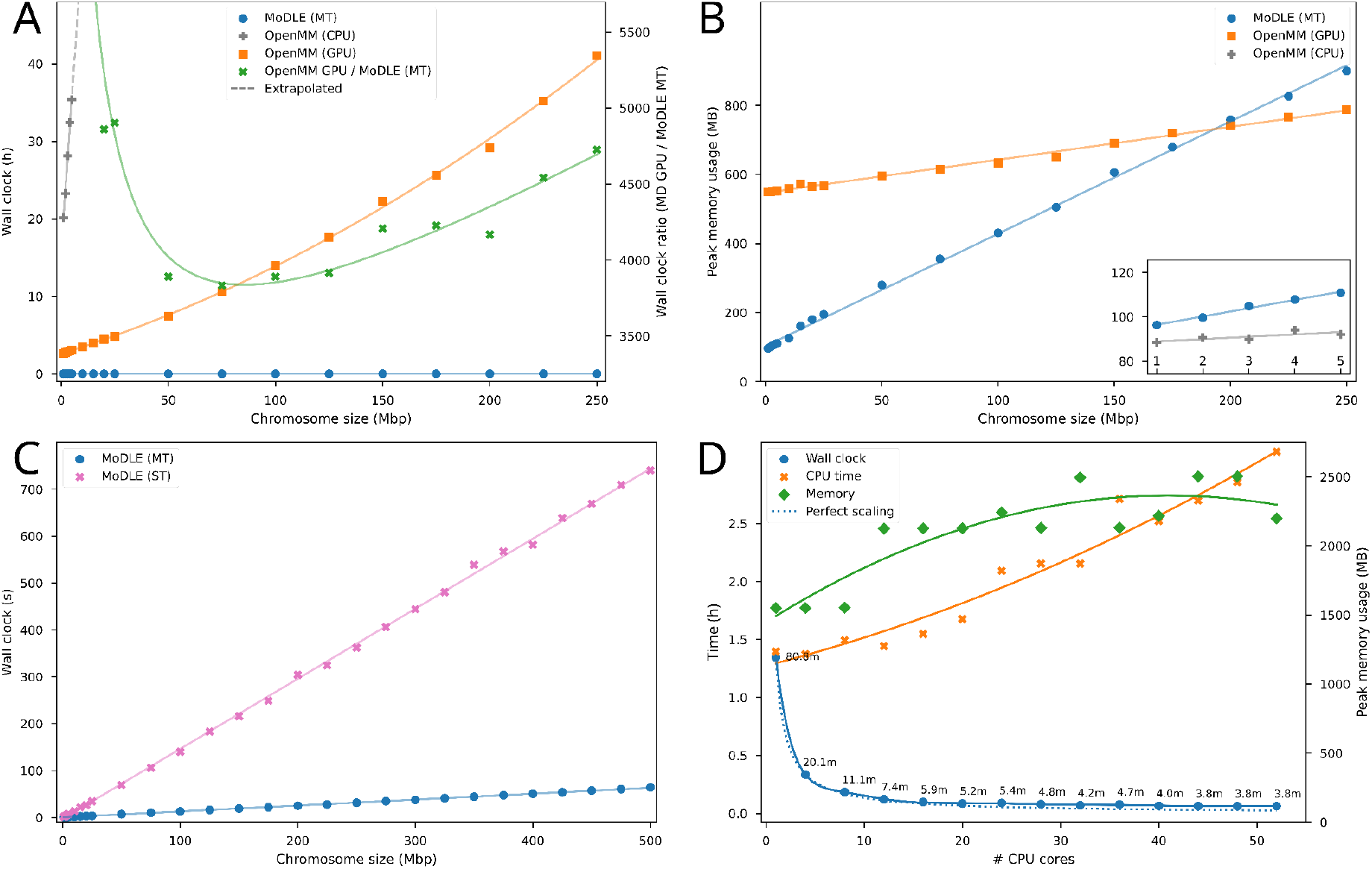
Benchmarking MoDLE and OpenMM. **A**: Median memory usage (in MBs) of MoDLE with multithreading (blue) compared to OpenMM with GPU (orange) for chromosome regions ranging in size from 1 to 250 Mbp. Inset shows comparison between MoDLE (blue) OpenMM with CPU (gray) for chromosome regions ranging in size from 1-5 Mbp. **B**: Median elapsed execution time (hours) of MoDLE with multithreading (blue), OpenMM with CPU (gray), OpenMM with GPU (orange) and the ratio of OpenMM (GPU) to MoDLE. Dotted lines are extrapolated. **C**: Comparison of the median elapsed execution time (seconds) of MoDLE with (blue) and without (pink) multithreading for chromosome regions ranging in size from 1 to 500 Mbp. **D**: Comparison of median elapsed execution time (hours) of MoDLE utilizing from 1-52 CPU cores. Blue line shows elapsed wall clock time (hours), whereas the orange line shows the CPU time (hours). The dotted line illustrates the corresponding theoretical perfect scaling of the executing time. Green line shows median peak memory usage (right axis; MB).

**Table 1:** Hardware specifications of computational resources used for simulation and benchmarking

| Identifier | CPU model | System memory | Operating system | Accelerator (GPU) |
| --- | --- | --- | --- | --- |
| Laptop A | Intel Core i9-9880H (8 cores) | 64 GB (4x 16 GB, DDR4 UDIMM 2667 MT/s dual-channel) | Arch Linux (Linux v5.17.1) | NVIDIA Quadro RTX 4000 (8 GB) |
| Server A | 2x AMD EPYC 7742 (2x 64 cores) | 2048 GB (32x 64 GB, RDIMM DDR4 2933 MT/s eight-channel) | RHEL 8.5 (Linux v4.18.0-305) | N/A |
| Server B | 2x Intel Xeon Gold 6138 (2x 20 cores) | 192 GB (12x 16 GB, RDIMM DDR4 2666 MT/s six-channel) | RHEL 7.9.2009 (Linux v3.10.0-1160.6.1) |  |
| Server C | 2x<br>Intel Xeon Gold 6230R<br>(2x 26 cores) | 192 GB (12x 16 GB,<br>RDIMM DDR4 2933<br>MT/s six-channel) |  |  |
| Server D | 2x<br>Intel Xeon Gold 6126<br>(2x 12 cores) | 384 GB (24x 16 GB,<br>RDIMM DDR4 2666<br>MT/s six-channel) |  | 4x<br>NVIDIA Tesla<br>P100 (16 GB) |

Comparing peak memory usage, MoDLE uses less memory than OpenMM for regions smaller than 200Mbp, and requires more memory for larger systems. Nevertheless, memory usage of both MoDLE and OpenMM scales linearly for increasing genome region sizes and is for all practical purposes within reasonable limits on today’s computers regardless of genome size (Fig. 3A-B).

Multithreading efficiently reduces MoDLE’s execution time for increasingly large genome sizes. With multithreading (52 CPU cores on server B; see Table 1), MoDLE can simulate loop extrusion contacts for a genome size of 500 Mbp in a little over one minute (Fig. 3C). Using a single thread (1 CPU core on server B; see Table 1), the same run takes around 12 minutes (Fig. 3C), which is still reasonable from a practical perspective and much faster than GPU accelerated OpenMM simulations. MoDLE peak memory usage is only slightly affected by multithreading, as each simulation instance only requires an additional 1-10 MB of memory (Fig. 3D). When simulating more than one chromosome, peak memory usage does not follow a simple linear pattern (Fig. 3D), as it is affected by the order in which simulation tasks are executed. This can lead to scenarios where for a brief period, two or more contact matrices are stored in system memory. We conclude that MoDLE, in contrast to OpenMM, runs efficiently even on systems with few CPU cores, such as laptop computers.

Further, we analyzed the strong scaling properties of MoDLE by simulating loop extrusion on the entire human genome (GRCh38; 3088Mbp). Increasing the number of CPU cores from 1 to 52, MoDLE execution time scales close to theoretical optimum (see Methods for details) (Fig. 3D; blue lines). Simulating loop extrusion on the human genome takes from 1 hour and 21 minutes (1 CPU core on server B; see Table 1) to 1 minute and 48 seconds (52 CPU cores on server B; see Table 1). We conclude that MoDLE can efficiently run on machines with a wide range of capabilities, ranging from laptop computers with 4-8 CPU cores, to multi-socket servers with over 50 CPU cores. Memory usage increases with the number of CPU cores, but never beyond reasonable limits on modern computers (Fig. 3D; orange line).

In conclusion, MoDLE is orders of magnitude faster than OpenMM in simulating loop extrusion contacts, and is especially efficient in simulating large genome regions or large input data sets. MoDLE can run efficiently on machines ranging from low-powered laptop computers to powerful multi-socket servers.

### Genome wide parameter optimization

Since MoDLE simulates genome-wide loop extrusion in a few minutes, systematic exploration of features underlying loop extrusion becomes feasible. To illustrate this point, we optimized the parameters underlying the modeled binding kinetics of CTCF. MoDLE implements this as a Markov process with an“Unbound” and a “Bound” state. With this model, the self-transition probabilities *P_UU_* and *P_BB_* specify how stably associated CTCF is once bound to DNA. The stationary distribution of the Markov chain reflects the probability of a given CTCF binding site to be bound (π_*B*_) in a simulation epoch (see Fig 4A). Simulation of *B* loop extrusion contacts using MoDLE or OpenMM can take advantage of ChIP-seq data from CTCF or cohesin to infer CTCF binding probabilities. Yet, when ChIP-seq data is not available it is possible to simulate loop extrusion using a constant and uniform CTCF binding probability that is chosen to optimize similarity with the Micro-C (or Hi-C) data. To optimize these parameters, we make use of an approach based on Bayesian optimization using Gaussian processes (see Methods). This optimization procedure attempts to minimize an objective function without making assumptions on its analytic form. To assess MoDLE’s performance we devised an objective function representing the similarity in stripe position and length between two contact matrices using H1-hESC Micro-C data (see Methods for details). After convergence (Fig. 4B), the optimization procedure revealed a range of near-optimal combinations of transition probabilities and CTCF occupancy probabilities instead of a single, optimal combination (Fig. 4C). Comparing the resulting loop contacts of selected parameter combinations with the optimal combination (π_*B*_ = 0. 747 and *P_UU_* = 0.963) confirms that CTCF can occupy its motif instances with probabilities ranging widely between 0.6 - 0.9 as long as the the stability of binding (*P_UU_*) is high (> 0.8). However, low binding stabilites (*P_UU_* < 0. 8) can also yield near-optimal concordance with the Micro-C data when CTCF occupancies > 0.9. Notably, the latter parameter combination is compatible with a dynamic exchange model where CTCF transiently occupies its motif instances, but still maintains stable loops [42]. From a selected set of parameter combinations (Fig. 4C), we simulated genome-wide loop extrusion contacts aiming at comparing these with Hi-C and Micro-C data. The resulting comparison shows that even uniform, optimized CTCF binding probabilities (Fig. 4C red star) can recapitulate many of the features seen in Micro-C and Hi-C data (Fig. 4D). Visualization of simulated contacts using a near-optimal parameter combination from another part of the plot (Fig. 4C; orange pentagon) reinforces that a range of parameter combinations can recapitulate the patterns seen in the Hi-C and Micro-C data (Fig. 4D). Selecting a suboptimal or non-optimal combination of parameters (Fig. 4C, green triangle and blue square) results in unrealistic contact patterns (Fig. 4D; see Supplementary Fig. 11 for an extensive comparison of different parameter combinations). In conclusion, MoDLE opens up for efficient exploration of parameters underlying DNA-DNA contact dynamics genome wide.

**Fig. 4:**
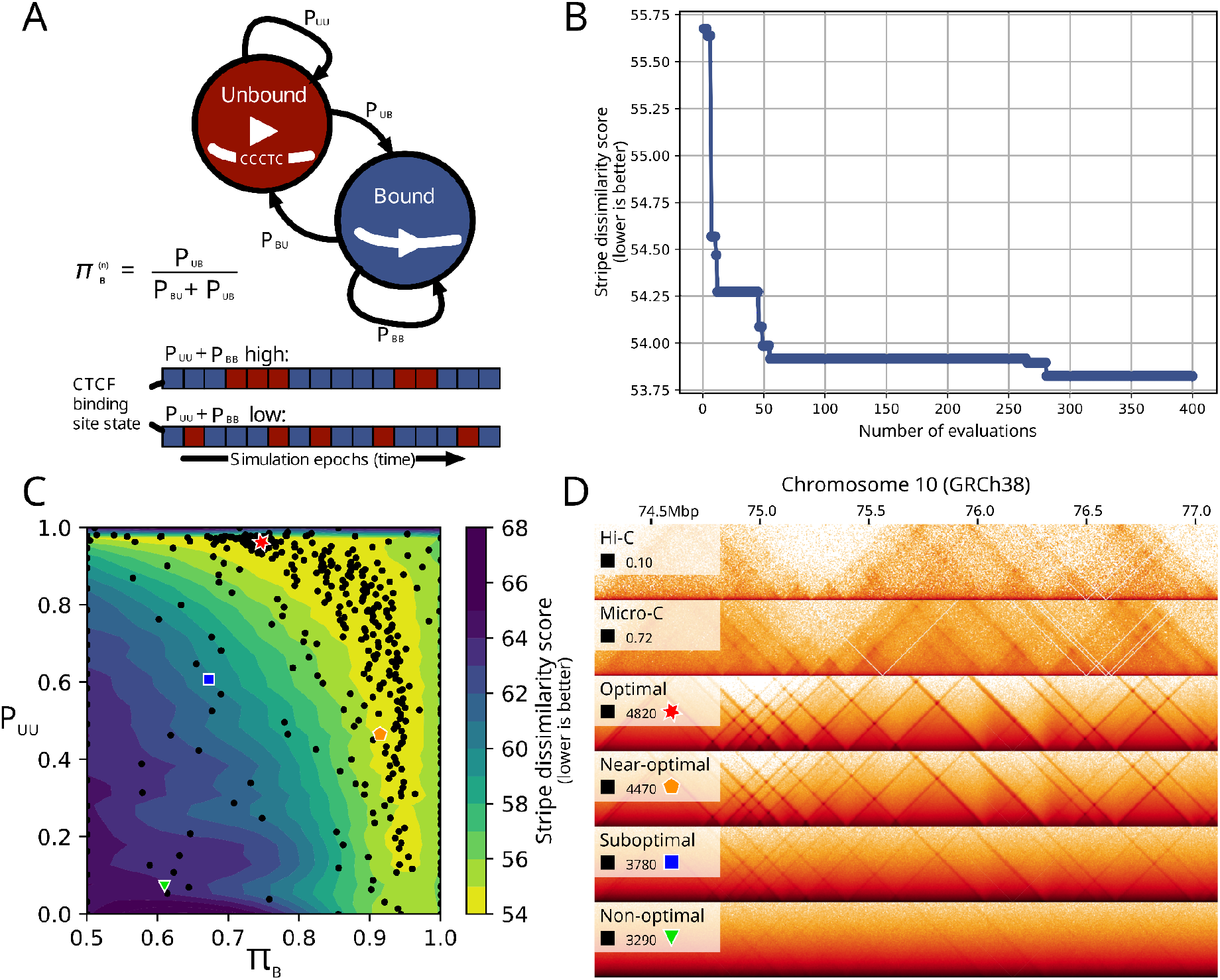
Genome-wide optimization of CTCF binding kinetics underlying loop extrusion. **A**: A Markov chain with an Unbound (red) and Bound (blue) state underlies MoDLE loop extrusion barrier modeling. The self-transition probability for the Bound state (*P_UU_*) reflects how stably barrier elements (i.e. CTCF) are bound to their binding sites. The stationary distribution of the Markov chain (π_B_) provides the CTCF binding probability at a given epoch in the simulation. The bottom diagram (red/blue boxes) shows an illustration of how the binding state (Bound in blue, and Unbound in red) of a single CTCF site would change during a simulation depending on *P_UU_* and *P_BB_*. **B**: Convergence of the objective function during the Bayesian optimization procedure. The objective function is a dissimilarity score comparing the pixels showing stripes and dots in the observed Micro-C data with the corresponding stripes and dots in the MoDLE output. See Methods (part 6) for details. C: Comparison of objective function in the parameter search space of *P_UU_* and π_*B*_. Optimal, near-optimal, suboptimal and non-optimal combinations are highlighted with a red star, orange pentagon, blue square and green triangle respectively. D: Side-by-side comparison of H1-hESC Micro-C data (top panel) and progressively less optimal combinations of *P_UU_* and π_*B*_ parameters.

### Predicting effects of TAD border alterations

To illustrate how MoDLE can be used to predict the effects of alterations to borders between TADs, we picked the well-characterized HoxD cluster which harbors several coordinated chromatin looping changes critical for proper limb formation in tetrapods [43,44]. We focused on deletions between the centromeric and telomeric domain (C-Dom and T-Dom, respectively) known to cause an increase in interactions between the two domains, including a rewiring of multiple enhancers. First, using the same parameter optimization approach described above, we inferred CTCF barrier occupancies in the wildtype condition based on JM8.N4 data. Then, we inactivated *(in silico)* inter-domain barrier elements by setting the occupancy of the CTCF motif instances to 0, and used MoDLE to simulate the resulting changes to the predicted loop extrusion contact maps. MoDLE correctly predicts that loops protrude beyond the deleted borders merging the two (C-Dom and T-Dom) TADs (Fig. 5). We also confirm that the border is highly resilient and requires a deletion of a large region encompassing the entire HoxD cluster to merge the TADs (see Fig. 5D-E). Inspecting enhancer signals in the region (Fig. 5E upper panel) confirms that the merging of the two domains indeed involves a rewiring of interactions of several enhancer elements, and a depletion of stripes at their borders. In conclusion, MoDLE can be used to predict changes to loop extrusion contact patterns from *in silico* alterations of TAD border properties.

**Fig. 5:**
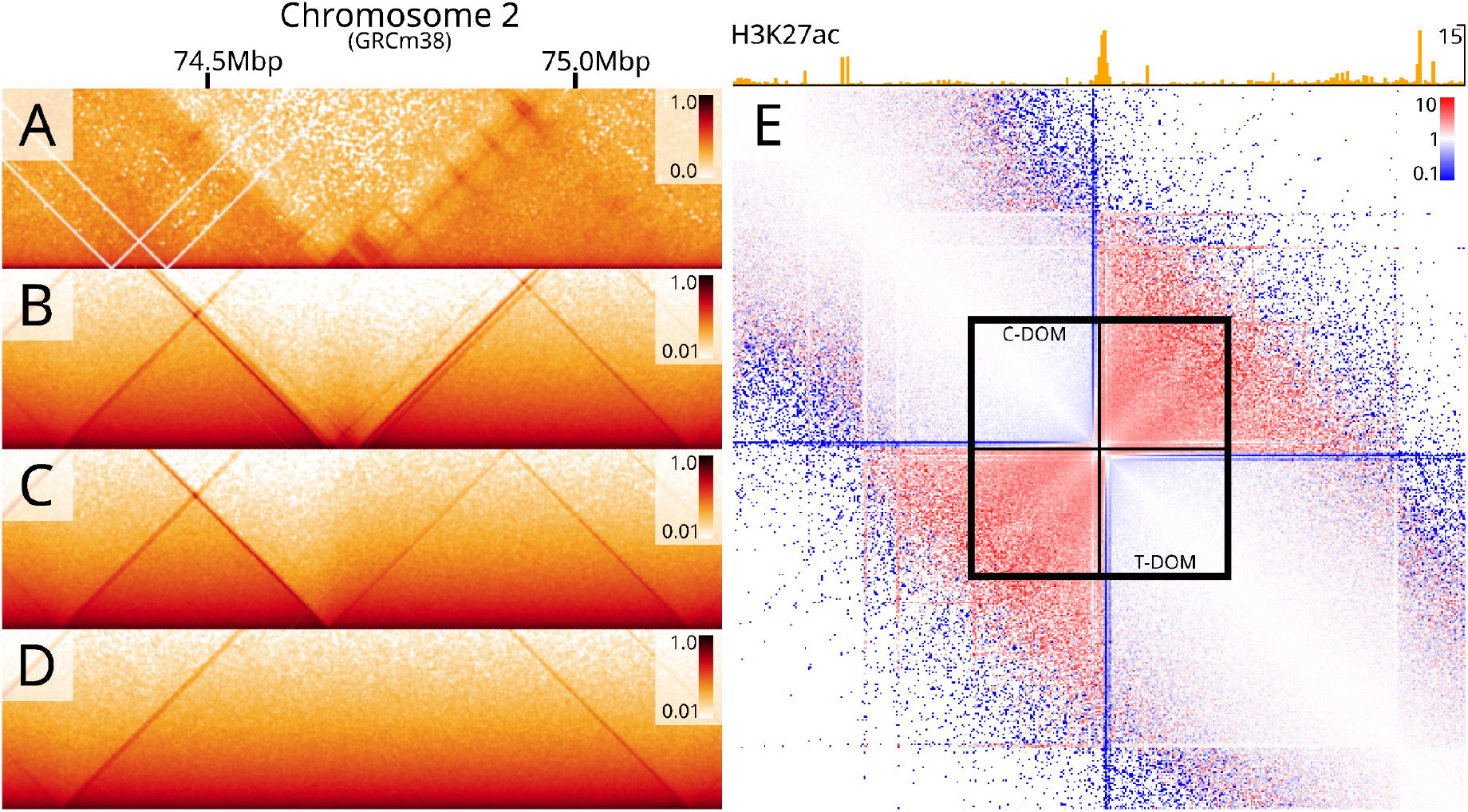
Using MoDLE to predict effects of deletions to TAD borders in the HoxD locus. **A**: Micro-C data in JM8.N4 mESC WT cells showing the interactions surrounding the HoxD cluster and the centromeric (C-DOM) and telomeric (T-DOM) domains in a non-mutated wildtype (WT) condition. B: MoDLE output from the same region in the WT condition. C: MoDLE output produced with a partial deletion of the border between the domains, D: MoDLE output with a complete deletion of the border between the domains. E: Differential contact map showing the ratio of MoDLE (WT condition; panel B) vs. MoDLE (full deletion, panel D). Regions enriched in MoDLE full deletion are shown in red, whereas regions enriched in MoDLE WT are shown in blue.

### Optimization of individual barrier parameters

In the absence of CTCF or Cohesin ChIP-seq data, MoDLE can utilize Micro-C or Hi-C data in combination with CTCF motif instances to effectively infer the occupancy of each individual barrier. To illustrate this, we selected a 5Mb region on chromosome 1 with 2103 CTCF candidate binding sites, corresponding to over 4000 parameters to be inferred. The large number of parameters for this genome region renders a Gaussian optimization approach computationally infeasible and inadequate. Thus, we developed a system to optimize extrusion barrier parameters using genetic algorithms (GA) (see Methods part 10 for details). A comparison of the input Micro-C data (Fig. 6A) and the corresponding optimized MoDLE output (Fig. 6B) shows that even without ChIP-seq information, MoDLE can be used to infer CTCF barrier occupancies individually to reproduce patterns seen in the Micro-C data. Comparing this MoDLE output with the corresponding output from MoDLE based on Rad21 ChIP-seq data (Fig. 6C) shows that TADs and borders are placed in analogous regions, yet with local differences in barrier strengths and stripe lengths. From MoDLE data simulated using optimized barrier occupancies (Fig. 6D) it is possible to compute the modeled binding profile of the LEF during the simulation (Fig. 6E; see section 9, Additional file 1 for details). Comparing these with ChIP-seq profiles of CTCF and Rad21 (Fig. 6F and G, respectively) shows that peaks and valleys coincide in a large fraction of regions, signifying that MoDLE can indeed infer biologically meaningful signals from its input data. We conclude that MoDLE, in the absence of ChIP-seq input data, can reliably infer CTCF occupancies of individual barriers to simulate loop extrusion contact patterns and to recapitulate binding profiles of CTCF and cohesin ChIP-seq data.

**Fig 6:**
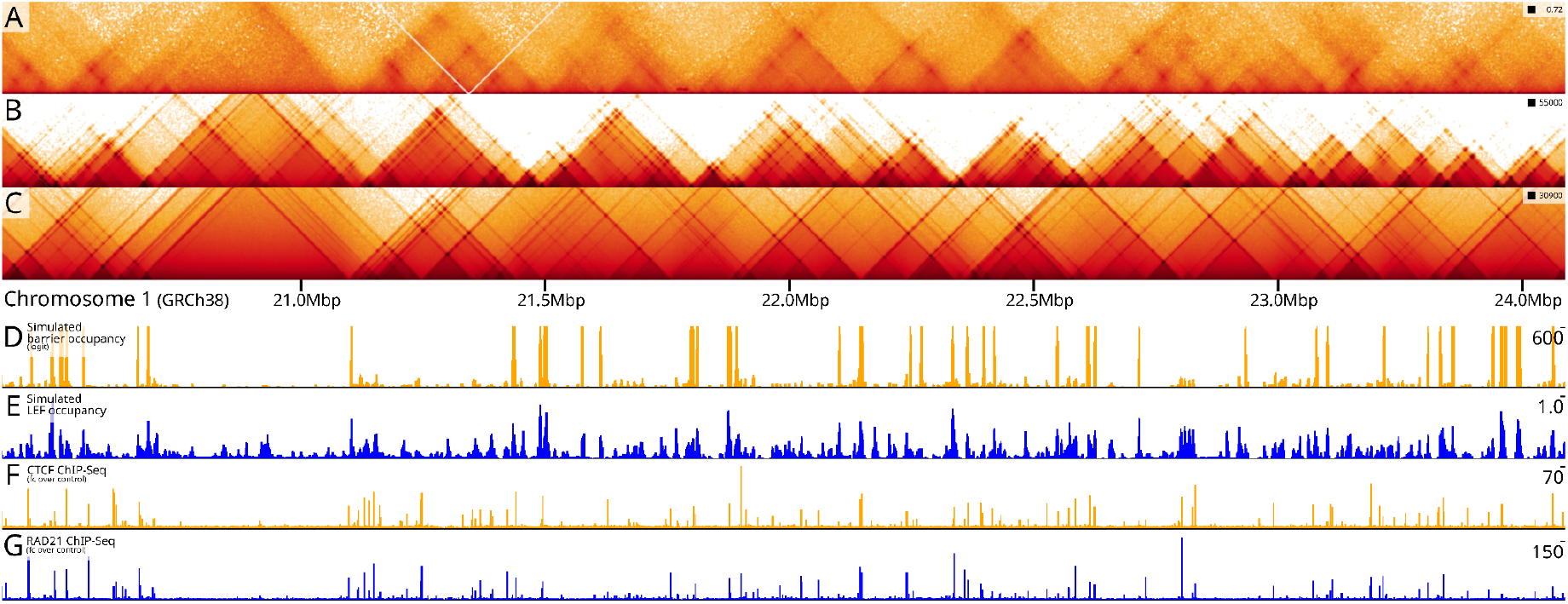
Optimization of individual barriers and computation of barrier and LEF profiles. **A**: Micro-C (hESC) data from a 5Mb region on chromosome 1 (20-25Mbp). **B**: MoDLE output for the same region, where individual barriers are optimized from Micro-C data. **C**: MoDLE output for the same region using Rad21 ChIP-seq data as input, **D**: Computed barrier occupancy profile from MoDLE trained on Micro-C data (normalized with *P_UU_* = 0.7), **E**: Computed LEF occupancy profile from MoDLE trained on Micro-C data. **F**: CTCF ChIP-seq data from the same region, **G**: Rad21 ChIP-seq data from the same region.

## Discussion

Efficient and realistic simulation of DNA-DNA spatial contacts is increasingly required for modeling and exploring genome structure and regulation. For example, our ability to reliably predict effects of mutations to TAD borders relies on available tools for simulating and comparing spatial contact data from normal and pathogenic states [14]. Further, simulations can be invaluable for exploring general genome folding principles [11] or underlying principles of loop extrusion [12,35,36]. Efficient tools for loop extrusion simulation will contribute to increasing our understanding of mechanisms ranging from gene regulation [1,2], to DNA repair [3]. MoDLE represents, to the best of our knowledge, the first command-line tool for high-throughput loop extrusion contact simulation. We expect MoDLE to supplement, rather than replace existing MD tools; especially in cases where large genome regions or large data sets need to be analyzed or simulated. This would in particular be the case for large-scale exploration of parameters underlying genome structure properties, as exemplified here for the binding kinetics of CTCF. In cases where Hi-C data is not available, we expect MoDLE to be useful for high-throughput loop extrusion contact prediction based on ChIP-seq, ATAC-seq or similar data in combination with CTCF motif instances (as exemplified in Fig. 2 and 4). In such cases, MoDLE could be useful for prediction of enhancer-promoter contacts aiding identification of functional regulatory interactions [45]. When Hi-C (or similar) data is available in a wildtype condition, MoDLE can be used for large scale prediction of mutations or alterations to TAD borders (as shown in Fig. 5 and 6). This would be useful for prioritization of mutations in genome editing settings.

New developments in experimental techniques augmented by integrated computational modeling, will continue to shed light on new genome organization principles at a rapid pace [46]. With MoDLE’s focus on computational speed and its modular architecture, new developments and knowledge are expected to easily be integrated into the tool to increase the complexity and realism of the underlying modeling parameters.

## Conclusions

We have developed MoDLE (Modeling of DNA Loop Extrusion), allowing high-performance stochastic modeling of DNA loop extrusion. MoDLE simulates loop extrusion contact matrices on large genome regions in a few minutes, even on low-powered laptop computers. MoDLE is available as a command line tool and can be accessed at github.com/paulsengroup/modle.

## Methods

### 1. MoDLE implementation and design overview

MoDLE is implemented in C++17 and is compiled with CMake. MoDLE uses a producer-consumer architecture where a single producer (a thread) communicates with multiple consumers through asynchronous message passing. The producer thread is responsible for reading input files and generating a set of simulation tasks to be consumed by a pool of worker threads. Tasks are implemented as light-weight C++ structs that are computationally cheap to generate and consume. A single task contains all the information needed for simulating DNA loop extrusion on a single chromosome in a specific simulation instance. Simulation instances are for the most part independent from each other and can thus run in parallel. We designed MoDLE such that each simulation instance requires less than 10 MB of memory to simulate loop extrusion on large mammalian chromosomes, such as chromosome 1 from the human genome. The space complexity of the thread–local state is linear with respect to the number of LEFs or extrusion barriers, whichever is largest. For a more detailed overview of MoDLE’s implementation see Section 1, Additional file 1.

Most of MoDLE’s memory budget is used to store molecular contacts generated by loop extrusion. MoDLE stores one instance of its custom contact matrix data structure for each chromosome that is being actively simulated. The space complexity of a contact matrix instance depends on the chromosome length, diagonal width and bin size. With default settings, representing contacts for chromosome 1 of the human genome requires approximately 120 MB of memory. Common operations on the contact matrix class are made thread-safe using lock striping implemented through hashing. For more details regarding the specialized contact matrix data structure refer to Section 2, Additional file 1. To achieve high-performance, MoDLE stores most of its data in contiguous memory using simple data structures such as vectors and bitsets (see Section 3, Additional file 1). Data is indexed such that extrusion barriers and extrusion units in a simulation instance can be efficiently traversed in 5’-3’ and 3’-5’ directions (see Section 8, Additional file 1). This allows MoDLE to bind/release LEFs, process collisions, register contacts and extrude DNA in linear time-complexity and with good locality of reference. The only step relying on an algorithm with super-linear time complexity is indexing, which has a worst-case time complexity of *O(n log n)* while approaching *O(n)* for the typical case.

More design and implementation details are available in Additional file 1. The latest MoDLE source code can be obtained in MoDLE’s GitHub repository: github.com/paulsengroup/modle

### 2. Running a simulation instance

The entire simulation instance is executed by a single worker thread and consists of the following phases:

- Wait until one or more tasks are available on the task queue.
- Setup the simulation internal state based on the task specification, this includes seeding the PRNG and setting the initial state for the extrusion barriers based on the occupancy (see Sections 1,3 and 4, Additional file 1).
- Run the simulation loop until a stopping criterion is met.

A single simulation epoch is articulated in the following steps:

- Select (inactive) LEFs that are currently not associated with DNA, and activate them. This is done by loading LEFs to a random position on the chromosome that is being simulated. The position is sampled from a uniform distribution (see Section 5, Additional file 1).
- Index extrusion units moving in the same direction so that they can be visited in 5’-3’ and 3’-5’ order (see Section 8, Additional file 1).
- Randomly select a subset of the active LEFs and use their position along the chromosome to generate molecular contacts in the chromosome contact matrix (see Section 9, Additional file 1).
- Generate candidate moves for each extrusion unit (see Section 10, Additional file 1).
- Update the extrusion barrier states by computing the next state in the Markov chain used to model extrusion barriers (see Section 6, Additional file 1).
- Detect collision events between LEFs and extrusion barriers as well as between LEFs (see Sections 12b-d, and g, Additional file 1).
- Update the candidate moves for extrusion units involved in collision events to satisfy the constraints imposed by the collision events (see Sections 12e-g, Additional file 1).
- Advance LEFs’ extrusion units by their respective moves (see Section 5, Additional file 1). Because of the preceding steps, this will yield a new valid simulation state, as moves have been updated to satisfy all the constraints imposed by collision events.
- Iterate over active LEFs and release them based on the outcome of a Bernoulli trial whose probability of success is computed based on the average LEF processivity and LEF state (e.g. LEFs whose extrusion units are involved in collision events with a pair of extrusion barriers in convergent orientation have a lower probability of being released). LEFs that are being released go back in the pool of available LEFs and will be loaded on a new genomic region during the next epoch (see Sections 5, Additional file 1).

MoDLE will continue iterating through the above steps until one of the simulation stopping criteria is met:

- A given number of epochs have been simulated
- Enough contacts have been registered to reach a target contact density. Both stopping criteria can be modified by users. By default, MoDLE will simulate loop extrusion until reaching an average contact density of 1 contact per pixel.

### 3. Hardware specifications

Analysis and benchmark code used to generate the data accompanying was run using the hardware specifications listed in Table 1.

### 4. MoDLE simulations

MoDLE’s data used for the heat map comparison shown in Fig. 2 were generated using the heatmap_comparison_pt1 Nextflow [47] workflow available on GitHub [48] and archived on Zenodo [49].

The list of candidate extrusion barrier positions and directions were generated by running MAST from the MEME suite [50] on GRCh38.p13 (GCF_000001405.39 [51] using the CTCF frequency matrix MA0139.1 from JASPAR 2022 [52].

The list of candidate barriers was then filtered using CTCF and RAD21 ChIP-seq data (fold-change over control and optimal IDR thresholded peaks). In brief, candidate barriers were intersected with the narrow-peak BED files for CTCF and RAD21. Then, each filtered barrier region was assigned with an occupancy computed by passing the RAD21 fold-change over control signal through a logistic function. Finally, the output of the logistic function was binned at 1 kbp to yield a barrier occupancy that is proportional to the number of CTCF motif instances as well as RAD21 fold-change over control signal. This procedure is largely based on [Fudenberg 2016]. The result of the procedure outlined above is a list of extrusion barrier occupancies binned at 1 kbp resolution. CTCF and RAD21 ChIP-seq for H1-hESC data was downloaded from ENCODE [53,54] (ENCFF255FRL [55], ENCFF473IZV [56], ENCFF821AQO [57] and ENCFF913JGA [58].

Contact matrices were generated using MoDLE v1.0.0-rc.7 with the parameters from Supplementary Table 1, Additional file 3. Parameters not listed in the table were left at default.

Contact matrices produced by MoDLE were then subsampled to an average contact density of 3 using cooltools random-sample v0.5.1 [59]. The resulting cooler files were then converted to multi-resolution cooler files using cooler zoomify [40]. Finally, multi-resolution contact matrices were visualized in HiGlass (v1.11.7) [60].

### 5. Molecular dynamics (OpenMM) simulations

Molecular dynamics data used for the heat map comparison in Fig. 2 were generated using the heatmap_comparison_pt1 Nextflow workflow available on GitHub [48] and archived on Zenodo [49]. This workflow uses OpenMM [37] to run MD simulations.

Simulation code is largely based on [61]. Simulations were carried out on 10 Mbp regions from chromosomes 2, 3, 5, 7 and 17 using a monomer size of 1 kbp and 200 kbp for LEF processivity and separation. Extrusion barrier positions, directions and occupancy were generated following the procedure outlined in Methods (part 1).

Contact matrices were generated with Polychrom [62] using a bin size of 5 kbp. The resulting cooler files were then converted to multi-resolution cooler files using cooler zoomify v0.8.11 [40].

### 6. Assessing loop extrusion feature similarities from contact frequencies

To objectively compare the contact matrices produced by MoDLE with contact matrices generated from Micro-C experiments and MD simulations we developed a specialized scoring algorithm. The algorithm was inspired by Stripenn [63].

The score is computed on rows and columns of a pair of contact matrices of identical resolutions transformed as follows.

First, both matrices are convolved using the difference of Gaussian (DoG). This highlights stripe and dot patterns found in contact matrices. Next, the transformed contact matrices are discretized using a step function mapping values below a certain threshold to 0 and all the others to 1. This results in two binary matrices, where non-zero pixels can be interpreted as part of a stripe or dot. Finally, we take advantage of the fact that stripes produced by loop extrusion always should start from the matrix diagonal. Thus, given a row or column of pixels starting on the matrix diagonal, and extending away from it, we stipulate that the last non-zero pixel in the vector of values represents the end of a stripe produced by DNA loop extrusion.

Given the above, we can measure the similarity of stripes between two contact matrices by considering the same row of pixels in a pair of contact matrices, computing the last non-zero pixels in both rows, and counting the number of matches. The same approach can be applied to columns of pixels. Finally, counting mismatches instead of matches can be used as a measure of dissimilarity. Contact matrix convolution and discretization, as well as computing this special score can be done using MoDLE’s helper tools (modle_tools transform and modle_tools evaluate respectively).

### 7. Contact matrix comparison

For comparison with MoDLE and OpenMM output, we used available Hi-C and Micro-C data from H1-hESC because these were of high resolution and had accompanying ChIP-seq data for both CTCF and RAD21 (4DNFIFJH2524 [64], 4DNFI9GMP2J8 [65], ENCFF255FRL [55], ENCFF473IZV [56], ENCFF821AQO [57] and ENCFF913JGA [58]). To assess stripe similarity of a pair of contact matrices we used the scoring algorithm described in Methods (part 6). The score was computed using Micro-C data as the ground truth. Pixel accuracy was computed as the ratio of correctly classified pixels to the total number of pixels in a 3 Mbp subdiagonal window around each barrier. The Pearson correlation between OpenMM and MoDLE was calculated based on all corresponding 5 kbp-pixel values in the 3 Mbp subdiagonal window of the OpenMM simulation regions.

### 8. Benchmark methodology

Benchmarks were run on a computing cluster using the run_benchmarks Nextflow workflow available on GitHub [48] and archived on Zenodo [49].

We ran two suites of benchmarks to assess the performance of MoDLE and compare it with that of molecular dynamics simulations based on OpenMM.

The first suite (Fig. 3A-C) compared the performance of MoDLE and OpenMM when simulating loop extrusion on an artificial chromosome with increasing length (ranging from 1 to 250 Mbp).

This benchmark was run using MoDLE (1 and 52 CPU cores) as well as OpenMM GPU and CPU implementation (1 CPU core, 1 GPU and 8 CPU cores respectively). CPU benchmarks were run on server C while benchmarks relying on GPU acceleration were run on server D (see Table 1). For OpenMM CPU implementation we limited the number of CPU cores to 8 (16 SMT cores) as the CPU implementation is known to not scale well past 16 threads [66]. OpenMM CPU implementation was used to simulate chromosome lengths up to 5 Mbp for practical reasons. MoDLE was run with default settings except for the number of cells, which was set to 104 to match the maximum number of available SMT cores.

OpenMM simulations were run using a monomer size of 2 kbp and LEF processivity and separation of 200 kbp.

The second suite of benchmarks involved simulating loop extrusion on the human genome (GRCh38) using MoDLE with a number of CPU cores ranging from 1 to 52. MoDLE was run with default settings except for the number of cells which was set to 104. The extrusion barrier annotation was generated as described in Methods (part 1).

In both cases, measurements were repeated 10 times for MoDLE and 5 times for OpenMM.

### 9. Genome-wide extrusion barrier parameter optimization

The genome-wide optimization of parameters affecting extrusion barrier occupancies was carried out using the gw_param_optimization Nextflow workflow available on GitHub [48] and archived on Zenodo [49].

The first step in the optimization procedure is running Stripenn v1.1.65.7 [63] on the H1-hESC Micro-C (4DNFI9GMP2J8 [67]) dataset to identify architectural stripes, which resulted in the identification of 5254 stripes. A small subset of these stripes were visually validated by comparing the annotated stripes with stripes that are visible from Micro-C data. Annotated stripes were split into two equally sized datasets by random sampling without replacement. One dataset was used for parameter optimization while the other was used for validation.

Parameter optimization is performed through the Bayesian optimization from scikit-optimize v0.9.0 [68] using an objective function based on the scoring metric described in Methods (part 6).

The parameters that are being optimized are the extrusion barrier occupancy (π_*B*_) and *P_UU_*, the self-transition probability of the unbound state.

The evaluation of the objective function proceeds as follows:

- A new genome-wide simulation is performed using the parameters proposed by the optimizer.
- The resulting cooler file is transformed with modle_tools transform by applying the difference of Gaussians followed by a binary discretization step, where pixel values above a certain threshold are discretized to 1 and all the others to 0.
- The score described in Methods (part 6) is then computed row and column-wise on the entire genome using modle_tools eval, producing two BigWig files. Here, the transformed Micro-C 4DNFI9GMP2J8 [67] dataset is used as reference.
- Scores are intersected with the extrusion barrier dataset for optimization and validation considering stripe direction (i.e. vertical stripes are intersected with column-wise scores while horizontal stripes are intersected with row-wise scores).
- Scores resulting from the intersection are then averaged, producing a floating-point number that is then returned to the optimizer, which will try to minimize this number.

In the transformation step a σ of 1.0 and 1.6 are used to generate the less and more blurry contact matrices to subtract when computing the difference of Gaussians. For the binary discretization of the Micro-C data a threshold of 1.5 was used, while simulated data was discretized using 0.75 as threshold.

The optimizer evaluated the objective function 400 times, each time computing the average score for the training and validation datasets.

Finally, the parameters that yielded the best score on the training dataset were used to generate a contact matrix in cooler format (see Fig. 4D, bottom panel).

### 10. Local extrusion barrier parameter optimization

The local extrusion barrier parameter optimization was carried out using the extrusion_barrier_param_optimization Nextflow workflow available on GitHub [48] and archived on Zenodo [49].

In brief, this workflow takes as input an extrusion barrier annotation consisting of barrier position and direction, and then optimizes the parameters for each individual barrier to maximize similarities with a reference HiC matrix.

The optimization approach is based on evolutionary algorithms (EAs) and was developed using primitives from the DEAP library [69].

Optimization was performed on a 5 Mbp region of the human chromosome 1 (20-25 Mbp, GRCh38) using the list of candidate CTCF binding sites overlapping this region as extrusion barrier annotation, for a total of 2103 extrusion barriers. Candidate CTCF binding sites were annotated using MAST as described in Methods (part 4). The H1-hESC Micro-C (4DNFI9GMP2J8 [70]) matrix was used as reference.

At a high level, the optimization workflow consists of running the same optimization script three times, using the output of an optimization run as input for the next run. The first run is tuned to favor exploration over exploitation, while the second and third runs used more conservative optimization parameters to progressively reduce the rate of exploration and favor exploitation.

The following is an overview of how the optimization strategy was developed:

- The optimization uses μ, λ as evolution strategy, where μ is the population size and λ is the number of offspring produced each generation. With this strategy, offspring that make it through the selection stage replace the previous population entirely. By default μ = 256 and λ = 512.
- Individuals are represented as two lists of real numbers of size *N,* where *N* is the number of extrusion barriers to be optimized. The first list of numbers represent s extrusion barrier occupancies (π_*B*_) while the second list represents the self-transition probability of the unbound state (*P_UU_*).
- Individuals are mutated by adding a relatively small offsets to 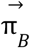 and 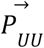. Offsets are drawn from a normal distribution with μ = 0 and σ set based on the desired degree of exploration. Values are clamped between 0.0 and 1.0, so mutating an individual always leads to another valid individual.
- The two-point crossover operator is used for mating.
- During selection, offsprings are sorted based on their fitness, and the top μ offsprings are selected to proceed to the next generation.
- The population is initialized differently depending on whether results from a previous optimization run are available.
  - Results from previous optimization are available: population initialized through random sampling with replacement from the set of fittest individuals that ever lived in the previous optimization run.
  - Otherwise, population is randomly initialized by generating μ individuals with π_*B*_ and P_*UU*_ set to random numbers drawn from the uniform distribution U(0.0, 1.0).
- Fitness is computed using a slightly modified version of the scoring function 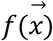 described in Methods (part 6). Function 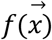 is not effective at guiding the optimization when occupancy is relatively low (e.g. < 0.5) and there are no stripes or dots in the reference matrix, as any parameter combination resulting in such a low occupancy produces no visible stripe or dot. To this end, we define a penalty function *p*(π_*B*_) that returns a coefficient between 1.0 and 2.0. The returned coefficient is close to 2.0 when π_*B*_ approaches 0.5, and rapidly falls to 1.0 when π_*B*_ moves towards 0.0 or 1.0. π_*B*_ very close to 1.0 are also penalized. See Supplementary Fig. 12 for more details regarding the penalty function *p*(π_*B*_). The output of the scoring function 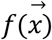 and penalty function *p*(π_B_) are multiplied together to produce the score used to compute the fitness of an individual 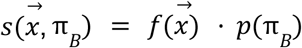. The fitness of an individual is computed as the average of the scores 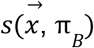 computed in correspondence of each extrusion barrier object of the optimization.
- The optimization runs until one of the following conditions is met:

- A target number of generations have been simulated (i.e. 1000 generations).
- Optimizer failed to significantly improve the population fitness (e.g. less than 1% fitness improvement over the last 25 generations).
- The population variability approaches 0.

To improve the performance of the optimizer on these regions we split the population into mainland population and one or more insular populations, and change some aspects of the optimization strategy.

First we initialize and optimize the mainland population (μ = 256 and λ = 512). When one of the stopping criteria is met, the fittest individuals from mainland are used to initialize the population of *m* islands. For each island, we randomly select and mask *k* consecutive alleles or barriers. *k* is generated by rounding a number drawn from a normal distribution with μ = 25 and σ = 5. 0. Crucially, masked barriers are inactive and are not allowed to mutate. For one of the *m* islands, instead of masking a random stretch of extrusion barriers, we inactivate all weak barriers when initializing the population. Thus, we replace alleles with π_*B*_ < 0.5 with the π_*B*_ = 0.0; *P_UU_* = 1. 0 allele. In this case, all loci are allowed to mutate.

Islands have μ = 128 and λ = 256. Islands evolve independently from each other and from the mainland, and follow the same stopping criteria used for the mainland.

Once all islands have been optimized, half of the mainland individuals are replaced with individuals from any of the islands. Island individuals are selected using fitness proportionate selection (i.e. random sampling with replacement, weighted by fitness).

Mainland and island optimization continue alternating until a total target number of mainland generations have been simulated, or when an optimization cycle fails to significantly improve the average mainland population fitness.

### 11. Simulations to predict the effect of TAD border alterations

Data for this section was generated using the comparison_with_mut Nextflow workflow available on GitHub [48] and archived on Zenodo [49].

Simulations were carried out using GRCm38.p6 as reference genome (GCF_000001635.26 [71].

CTCF and RAD21 ChIP-seq fold-change over control for JM8.N4 was generated by processing data from GSE90994 [72], (SRR5085152 [73], SRR5085153 [74], SRR5085154 [75], SRR5085155 [76], SRR5085156 [77], SRR5085157 [78] using the ENCODE ChIP-seq pipeline v2 [79] using ENCODE4 genomic datasets for GRCm38.

The wild-type extrusion barrier annotation was generated following the procedure outlined in Methods (part 4).

The barrier annotation was further refined using the parameter optimization strategy described in Methods (part 10) using a JM8.N4 Micro-C dataset as reference (4DNFINNZDDXV [80]).

The optimized extrusion barrier annotation was then mutated by removing extrusion barriers overlapping the del1-13d9lac and delattP-Rel5d9lac regions from Supplemental Figure S2 in Rodríguez-Carballo 2017 [43].

## Abbreviations

3D: Three-dimensional
BED: Browser Extensible Data
CLI: Command line interface
CTCF: CCCTC binding factor
hESC: Human Embryonic Stem Cells
kbp: kilo basepair
LEF: Loop Extruding Factor
Mbp: Mega basepair
MD: Molecular dynamics
PRNG: Pseudorandom Number Generator
SMC: Structural Maintenance of Chromosomes
SMT: Simultaneous Multithreading
TAD: Topologically Associating Domain

## Declarations

### Ethics approval and consent to participate

Not applicable

### Consent for publication

Not applicable

### Availability of data and materials

Project name: MoDLE

Project home page: https://github.com/paulsengroup/modle

Archived version: 10.5281/zenodo.6424697

License: MIT

Operative system(s): UNIX-like (Platform independent when using containers) Programming language: C++

Other requirements:

- C++17 compiler (e.g. GCC 8+, Clang8+, AppleClang 10+)
- CMake 3.18 or newer
- Conan 1.50 or newer
- Python3 (to install and run Conan)
- Scipy (Python3 package, required to run unit tests)
- wCorr (R package, required to run unit tests)

The complete list of MoDLE dependencies is available in Supplementary Table 2, Additional file 3. Dependency installation is automated using CMake and Conan.

MoDLE source code is available on GitHub at https://github.com/paulsengroup/modle [81] and is archived on Zenodo at https://doi.org/10.5281/zenodo.6424697 [82]. MoDLE’s version used throughout the manuscript is MoDLE v1.0.0-rc.7 [83], except for performance benchmarks which used MoDLE v1.0.0-rc.2 [84].

Code used for the data analysis is available on GitHub at https://github.com/paulsengroup/2021-modle-paper-001-data-analysis[48] and is archived on Zenodo at https://doi.org/10.5281/zenodo.7072939[49].

Data produced by running the runme.sh script from the data analysis core repository, including simulated contact matrices in cooler format were archived on Zenodo (DOI: 10.5281/zenodo.7074561).

Reference genome assembly and assembly report for GCF_000001405.39_GRCh38.p13 [51] and GCF_000001635.26 [85] were downloaded through the NCBI FTP server [86].

ChIP-seq data for the following accession numbers were downloaded from the ENCODE portal [53,54]: ENCFF255FRL [55], ENCFF473IZV [56], ENCFF821AQO [57] and ENCFF913JGA [58].

ChIP-seq data for the following accession number was downloaded from the Gene Expression Omnibus: GSM4665702 [87].

ChIP-seq sequencing data for GSE90994 [88] were downloaded from EBI’s mirror of the SRA: SRR5085152 [89], SRR5085153 [90], SRR5085154 [91], SRR5085155 [92], SRR5085156 [93], SRR5085157 [94].

H1-hESC Hi-C and Micro-C data, as well as JM8.N4 Micro-C data in multi-resolution cooler format (4DNFIFJH2524 [64], 4DNFI9GMP2J8 [67], 4DNFINNZDDXV [80] were downloaded from the 4DNucleome Data Portal [95].

The frequency matrix in MEME format for the CTCF motif (MA0139.1) was downloaded from JASPAR 2022 CORE non-redundant database [52].

### Authors’ contributions

JP and RR conceived and designed the study; RR developed the software and the analysis code; RR performed benchmarking analysis, genome wide and local parameter optimization as well as the comparative analysis for the HoxD mutants; JP performed comparison analyses of MoDLE with OpenMM and Micro-C data; JP and RR wrote the original draft; AM helped with the manuscript revising; VK, AM and TR provided critical feedback and assisted in the improvement of the tool and analyses; JP supervised the project. All authors read and approved the final manuscript.

## Acknowledgements

The simulations were performed on resources provided by Sigma2 - the National Infrastructure for High Performance Computing and Data Storage in Norway, with account number NN8041K.

We thank the ENCODE Consortium and the labs of Peter Park and Job Dekker for contributing with ENCODE data used as part of this study.

## Supplementary material

- Additional file 1 (pdf): Supplementary text. Detailed description of MoDLE’s underlying simulation model and implementation.
- Additional file 2 (pdf): Supplementary figures. Supplementary Figures 1-12.
- Additional file 3 (pdf): Supplementary tables. Supplementary Tables 1-2.

## References

1. Braccioli L, de Wit E. CTCF: a Swiss-army knife for genome organization and transcription regulation. Essays Biochem. 2019;63:157–65.

2. Razin SV, Gavrilov AA, Vassetzky YS, Ulianov SV. Topologically-associating domains: gene warehouses adapted to serve transcriptional regulation. Transcription. 2016;7:84–90.

3. Arnould C, Rocher V, Finoux A-L, Clouaire T, Li K, Zhou F, et al. Loop extrusion as a mechanism for formation of DNA damage repair foci. Nature. 2021;590:660–5.

4. Peters J-M. How DNA loop extrusion mediated by cohesin enables V(D)J recombination. Curr Opin Cell Biol. 2021;70:75–83.

5. Goloborodko A, Marko JF, Mirny LA. Chromosome Compaction by Active Loop Extrusion. Biophys J. 2016;110:2162–8.

6. Ganji M, Shaltiel IA, Bisht S, Kim E, Kalichava A, Haering CH, et al. Real-time imaging of DNA loop extrusion by condensin. Science. 2018;360:102–5.

7. Golfier S, Quail T, Kimura H, Brugués J. Cohesin and condensin extrude DNA loops in a cell cycle-dependent manner. Elife [Internet]. 2020;9. Available from: http://dx.doi.org/10.7554/eLife.53885

8. Lieberman-Aiden E, van Berkum NL, Williams L, Imakaev M, Ragoczy T, Telling A, et al. Comprehensive mapping of long-range interactions reveals folding principles of the human genome. Science. 2009;326:289–93.

9. Hsieh T-HS, Weiner A, Lajoie B, Dekker J, Friedman N, Rando OJ. Mapping Nucleosome Resolution Chromosome Folding in Yeast by Micro-C. Cell. 2015;162:108–19.

10. Krietenstein N, Abraham S, Venev SV, Abdennur N, Gibcus J, Hsieh T-HS, et al. Ultrastructural Details of Mammalian Chromosome Architecture. Mol Cell. 2020;78:554–65.e7.

11. Sanborn AL, Rao SSP, Huang S-C, Durand NC, Huntley MH, Jewett AI, et al. Chromatin extrusion explains key features of loop and domain formation in wild-type and engineered genomes. Proc Natl Acad Sci U S A. 2015;112:E6456–65.

12. Fudenberg G, Imakaev M, Lu C, Goloborodko A, Abdennur N, Mirny LA. Formation of Chromosomal Domains by Loop Extrusion. Cell Rep. 2016;15:2038–49.

13. Brandâo HB, Ren Z, Karaboja X, Mirny LA, Wang X. DNA-loop-extruding SMC complexes can traverse one another in vivo. Nat Struct Mol Biol. 2021;28:642–51.

14. Lupiáñez DG, Spielmann M, Mundlos S. Breaking TADs: How Alterations of Chromatin Domains Result in Disease. Trends Genet. 2016;32:225–37.

15. Alipour E, Marko JF. Self-organization of domain structures by DNA-loop-extruding enzymes. Nucleic Acids Res. 2012;40:11202–12.

16. Bauer BW, Davidson IF, Canena D, Wutz G, Tang W, Litos G, et al. Cohesin mediates DNA loop extrusion by a “swing and clamp” mechanism. Cell. 2021;184:5448–64.e22.

17. Golov AK, Golova AV, Gavrilov AA, Razin SV. Sensitivity of cohesin-chromatin association to high-salt treatment corroborates non-topological mode of loop extrusion. Epigenetics Chromatin. 2021;14:36.

18. Pradhan B, Barth R, Kim E, Davidson IF, Bauer B, van Laar T, et al. SMC complexes can traverse physical roadblocks bigger than their ring size [Internet]. bioRxiv. 2021 [cited 2022 Apr 11]. p. 2021.07.15.452501. Available from: https://www.biorxiv.org/content/10.1101/2021.07.15.452501v1.abstract

19. Rao SSP, Huntley MH, Durand NC, Stamenova EK, Bochkov ID, Robinson JT, et al. A 3D map of the human genome at kilobase resolution reveals principles of chromatin looping. Cell. 2014;159:1665–80.

20. Vian L, Pękowska A, Rao SSP, Kieffer-Kwon K-R, Jung S, Baranello L, et al. The Energetics and Physiological Impact of Cohesin Extrusion. Cell. 2018;173:1165–78.e20.

21. Tedeschi A, Wutz G, Huet S, Jaritz M, Wuensche A, Schirghuber E, et al. Wapl is an essential regulator of chromatin structure and chromosome segregation. Nature. 2013;501:564–8.

22. Haarhuis JHI, van der Weide RH, Blomen VA, Yáñez-Cuna JO, Amendola M, van Ruiten MS, et al. The Cohesin Release Factor WAPL Restricts Chromatin Loop Extension. Cell. 2017;169:693–707.e14.

23. Nakamura R, Motai Y, Kumagai M, Wike CL, Nishiyama H, Nakatani Y, et al. CTCF looping is established during gastrulation in medaka embryos. Genome Res. 2021;31:968–80.

24. Wutz G, Várnai C, Nagasaka K, Cisneros DA, Stocsits RR, Tang W, et al. Topologically associating domains and chromatin loops depend on cohesin and are regulated by CTCF, WAPL, and PDS5 proteins. EMBO J. 2017;36:3573–99.

25. Rao SSP, Huang S-C, Glenn St Hilaire B, Engreitz JM, Perez EM, Kieffer-Kwon K-R, et al. Cohesin Loss Eliminates All Loop Domains. Cell. 2017;171:305–20.e24.

26. Liu NQ, Magnitov M, Schijns M, van Schaik T, van der Weide RH, Teunissen H, et al. Rapid depletion of CTCF and cohesin proteins reveals dynamic features of chromosome architecture [Internet]. bioRxiv. 2021 [cited 2022 Apr 11]. p. 2021.08.27.457977. Available from: https://www.biorxiv.org/content/10.1101/2021.08.27.457977v1.full

27. Nora EP, Goloborodko A, Valton A-L, Gibcus JH, Uebersohn A, Abdennur N, et al. Targeted Degradation of CTCF Decouples Local Insulation of Chromosome Domains from Genomic Compartmentalization. Cell. 2017;169:930–44.e22.

28. Barbieri M, Chotalia M, Fraser J, Lavitas L-M, Dostie J, Pombo A, et al. Complexity of chromatin folding is captured by the strings and binders switch model. Proc Natl Acad Sci U S A. 2012;109:16173–8.

29. Naumova N, Imakaev M, Fudenberg G, Zhan Y, Lajoie BR, Mirny LA, et al. Organization of the mitotic chromosome. Science. 2013;342:948–53.

30. Jost D, Carrivain P, Cavalli G, Vaillant C. Modeling epigenome folding: formation and dynamics of topologically associated chromatin domains. Nucleic Acids Res. 2014;42:9553–61.

31. Benedetti F, Dorier J, Burnier Y, Stasiak A. Models that include supercoiling of topological domains reproduce several known features of interphase chromosomes. Nucleic Acids Res. 2014;42:2848–55.

32. Goloborodko A, Imakaev MV, Marko JF, Mirny L. Compaction and segregation of sister chromatids via active loop extrusion. Elife [Internet]. 2016;5. Available from: http://dx.doi.org/10.7554/eLife.14864

33. Anderson JA, Glaser J, Glotzer SC. HOOMD-blue: A Python package for high-performance molecular dynamics and hard particle Monte Carlo simulations. Comput Mater Sci. 2020;173:109363.

34. Schwarzer W, Abdennur N, Goloborodko A, Pekowska A, Fudenberg G, Loe-Mie Y, et al. Two independent modes of chromatin organization revealed by cohesin removal. Nature. 2017;551:51–6.

35. Nuebler J, Fudenberg G, Imakaev M, Abdennur N, Mirny LA. Chromatin organization by an interplay of loop extrusion and compartmental segregation. Proc Natl Acad Sci U S A. 2018;115:E6697–706.

36. Banigan EJ, van den Berg AA, Brandão HB, Marko JF, Mirny LA. Chromosome organization by one-sided and two-sided loop extrusion. Elife [Internet]. 2020;9. Available from: http://dx.doi.org/10.7554/eLife.53558

37. Eastman P, Swails J, Chodera JD, McGibbon RT, Zhao Y, Beauchamp KA, et al. OpenMM 7: Rapid development of high performance algorithms for molecular dynamics. PLoS Comput Biol. 2017;13:e1005659.

38. Buckle A, Brackley CA, Boyle S, Marenduzzo D, Gilbert N. Polymer Simulations of Heteromorphic Chromatin Predict the 3D Folding of Complex Genomic Loci. Mol Cell. 2018;72:786–97.e11.

39. Zhang S, Übelmesser N, Josipovic N, Forte G, Slotman JA, Chiang M, et al. RNA polymerase II is required for spatial chromatin reorganization following exit from mitosis. Sci Adv. 2021;7:eabg8205.

40. Abdennur N, Mirny LA. Cooler: scalable storage for Hi-C data and other genomically labeled arrays. Bioinformatics. 2020;36:311–6.

41. Gabriele M, Brandão HB, Grosse-Holz S, Jha A, Dailey GM, Cattoglio C, et al. Dynamics of CTCF and cohesin mediated chromatin looping revealed by live-cell imaging [Internet]. bioRxiv. 2021 [cited 2022 Apr 11]. p. 2021.12.12.472242. Available from: https://www.biorxiv.org/content/10.1101/2021.12.12.472242v1

42. Hansen AS, Pustova I, Cattoglio C, Tjian R, Darzacq X. CTCF and cohesin regulate chromatin loop stability with distinct dynamics. Elife [Internet]. 2017;6. Available from: http://dx.doi.org/10.7554/eLife.25776

43. Rodríguez-Carballo E, Lopez-Delisle L, Zhan Y, Fabre PJ, Beccari L, El-Idrissi I, et al. The cluster is a dynamic and resilient TAD boundary controlling the segregation of antagonistic regulatory landscapes. Genes Dev. 2017;31:2264–81.

44. Rodríguez-Carballo E, Lopez-Delisle L, Willemin A, Beccari L, Gitto S, Mascrez B, et al. Chromatin topology and the timing of enhancer function at the locus. Proc Natl Acad Sci U S A. 2020;117:31231–41.

45. Xu H, Zhang S, Yi X, Plewczynski D, Li MJ. Exploring 3D chromatin contacts in gene regulation: The evolution of approaches for the identification of functional enhancer-promoter interaction. Comput Struct Biotechnol J. 2020;18:558–70.

46. Di Stefano M, Paulsen J, Jost D, Marti-Renom MA. 4D nucleome modeling. Curr Opin Genet Dev. 2021;67:25–32.

47. Di Tommaso P, Chatzou M, Floden EW, Barja PP, Palumbo E, Notredame C. Nextflow enables reproducible computational workflows. Nat Biotechnol. 2017;35:316–9.

48. 2021-modle-paper-001-data-analysis: Data analysis code for the first paper about MoDLE (preprint available soon) [Internet]. Github; [cited 2022 Apr 11]. Available from: https://github.com/paulsengroup/2021-modle-paper-001-data-analysis

49. Rossini R, Kumar V, Mathelier A, Rognes T, Paulsen J. Data analysis code for: “MoDLE: High-performance stochastic modeling of DNA loop extrusion interactions.” 2022 [cited 2022 Sep 13]; Available from: https://zenodo.org/record/7072939

50. Bailey TL, Gribskov M. Combining evidence using p-values: application to sequence homology searches. Bioinformatics. 1998;14:48–54.

51. GRCh38.p13 - hg38 - Genome - Assembly - NCBI [Internet]. [cited 2022 Apr 12]. Available from: https://identifiers.org/assembly:GCF_000001405.39

52. Castro-Mondragon JA, Riudavets-Puig R, Rauluseviciute I, Lemma RB, Turchi L, Blanc-Mathieu R, et al. JASPAR 2022: the 9th release of the open-access database of transcription factor binding profiles. Nucleic Acids Res. 2022;50:D165–73.

53. ENCODE Project Consortium. An integrated encyclopedia of DNA elements in the human genome. Nature. 2012;489:57–74.

54. Davis CA, Hitz BC, Sloan CA, Chan ET, Davidson JM, Gabdank I, et al. The Encyclopedia of DNA elements (ENCODE): data portal update. Nucleic Acids Res. 2018;46:D794–801.

55. ENCFF255FRL – ENCODE [Internet]. [cited 2022 Apr 12]. Available from: https://identifiers.org/encode:ENCFF255FRL

56. ENCFF473IZV – ENCODE [Internet]. [cited 2022 Apr 12]. Available from: https://identifiers.org/encode:ENCFF473IZV

57. ENCFF821AQO – ENCODE [Internet]. [cited 2022 Apr 12]. Available from: https://identifiers.org/encode:ENCFF821AQO

58. Encff913jga – encode [Internet]. [cited 2022 Apr 12]. Available from: https://identifiers.org/encode:ENCFF913JGA

59. Venev S, Abdennur N, Goloborodko A, Flyamer I, Fudenberg G, Nuebler J, et al. open2c/cooltools: v0.5.1 [Internet]. 2022. Available from: https://zenodo.org/record/6324229

60. Kerpedjiev P, Abdennur N, Lekschas F, McCallum C, Dinkla K, Strobelt H, et al. HiGlass: web-based visual exploration and analysis of genome interaction maps. Genome Biol. 2018;19:125.

61. Banigan EJ, Mirny LA. The interplay between asymmetric and symmetric DNA loop extrusion. Elife [Internet]. 2020;9. Available from: http://dx.doi.org/10.7554/eLife.63528

62. Imakaev M, Goloborodko A, hbbrandao. mirnylab/polychrom: v0.1.0 [Internet]. 2019. Available from: https://zenodo.org/record/3579473

63. Yoon S, Chandra A, Vahedi G. Stripenn detects architectural stripes from chromatin conformation data using computer vision. Nat Commun. 2022;13:1602.

64. 4DNFIFJH2524.mcool – 4DN Data Portal [Internet]. [cited 2022 Sep 13]. Available from: https://identifiers.org/4dn:4DNFIFJH2524

65. 4DNFI9GMP2J8.mcool – 4DN Data Portal [Internet]. [cited 2022 Sep 13]. Available from: https://identifiers.org/4dn:4DNFI9GMP2J8

66. openmm [Internet]. Github; [cited 2022 Apr 11]. Available from: https://github.com/openmm/openmm/issues/3267

67. 4DNFI9GMP2J8.Mcool – 4DN data portal [Internet]. [cited 2022 Apr 12]. Available from: https://identifiers.org/4dn:4DNFI9GMP2J8

68. scikit-optimize: Sequential model-based optimization with a ‘scipy.optimize’ interface [Internet]. Github; [cited 2022 Apr 11]. Available from: https://github.com/scikit-optimize/scikit-optimize

69. Fortin F-A, De Rainville F-M, Gardner M-A, Parizeau M, Gagné C. DEAP: Evolutionary Algorithms Made Easy. J Mach Learn Res. 2012;13:2171–5.

70. 4DNFI9GMP2J8.mcool – 4DN Data Portal [Internet]. [cited 2022 Sep 13]. Available from: https://identifiers.org/4dn:4DNFI9GMP2J8

71. GRCm38.p6 - Genome - Assembly - NCBI [Internet]. [cited 2022 Sep 13]. Available from: https://identifiers.org/assembly:GCF_000001635.26

72. GEO Accession viewer [Internet]. [cited 2022 Sep 13]. Available from: https://identifiers.org/GEO:GSE90994

73. GSM2418858: WT_IgG_ChIPSeq; Mus musculus; ChIP-Seq - SRA - NCBI [Internet]. [cited 2022 Sep 13]. Available from: https://identifiers.org/insdc.sra:SRR5085152

74. GSM2418858: WT_IgG_ChIPSeq; Mus musculus; ChIP-Seq - SRA - NCBI [Internet]. [cited 2022 Sep 13]. Available from: https://identifiers.org/insdc.sra:SRR5085153

75. GSM2418859: WT_Rad21_ChIPSeq; Mus musculus; ChIP-Seq - SRA - NCBI [Internet]. [cited 2022 Sep 13]. Available from: https://identifiers.org/insdc.sra:SRR5085154

76. GSM2418859: WT_Rad21_ChIPSeq; Mus musculus; ChIP-Seq - SRA - NCBI [Internet]. [cited 2022 Sep 13]. Available from: https://identifiers.org/insdc.sra:SRR5085155

77. GSM2418860: WT_CTCF_ChIPSeq; Mus musculus; ChIP-Seq - SRA - NCBI [Internet]. [cited 2022 Sep 13]. Available from: https://identifiers.org/insdc.sra:SRR5085156

78. GSM2418860: WT_CTCF_ChIPSeq; Mus musculus; ChIP-Seq - SRA - NCBI [Internet]. [cited 2022 Sep 13]. Available from: https://identifiers.org/insdc.sra:SRR5085157

79. GitHub - ENCODE-DCC/chip-seq-pipeline2: ENCODE ChIP-seq pipeline [Internet]. GitHub. [cited 2022 Sep 13]. Available from: https://github.com/ENCODE-DCC/chip-seq-pipeline2

80. 4DNFINNZDDXV.mcool – 4DN Data Portal [Internet]. [cited 2022 Sep 13]. Available from: https://identifiers.org/4dn:4DNFINNZDDXV

81. modle: High-performance stochastic modeling of DNA loop extrusion interactions [Internet]. Github; [cited 2022 Apr 11]. Available from: https://github.com/paulsengroup/modle

82. Rossini R, Kumar V, Mathelier A, Rognes T, Paulsen J. MoDLE [Internet]. Zenodo; 2022. Available from: https://zenodo.org/record/6424697

83. Rossini R, Kumar V, Mathelier A, Rognes T, Paulsen J. MoDLE. 2022 [cited 2022 Sep 13]; Available from: https://zenodo.org/record/6992533

84. Rossini R, Vipin K, Mathelier A, Rognes T, Paulsen J. MoDLE: High-performance stochastic modeling of DNA loop extrusion interactions [Internet]. 2022. Available from: https://zenodo.org/record/6424890

85. GRCm38.p6 - Genome - Assembly - NCBI [Internet]. [cited 2022 Sep 13]. Available from: https://identifiers.org/assembly:GCF_000001635.26

86. O’Leary NA, Wright MW, Brister JR, Ciufo S, Haddad D, McVeigh R, et al. Reference sequence (RefSeq) database at NCBI: current status, taxonomic expansion, and functional annotation. Nucleic Acids Res. 2016;44:D733–45.

87. GEO Accession viewer [Internet]. [cited 2022 Sep 13]. Available from: https://identifiers.org/GEO:GsM4665702

88. GEO Accession viewer [Internet]. [cited 2022 Sep 13]. Available from: https://identifiers.org/GEO:GsE90994

89. GSM2418858: WT_IgG_ChIPSeq; Mus musculus; ChIP-Seq - SRA - NCBI [Internet]. [cited 2022 Sep 13]. Available from: https://identifiers.org/insdc.sra:SRR5085152

90. GSM2418858: WT_IgG_ChIPSeq; Mus musculus; ChIP-Seq - SRA - NCBI [Internet]. [cited 2022 Sep 13]. Available from: https://identifiers.org/insdc.sra:SRR5085153

91. GSM2418859: WT_Rad21_ChIPSeq; Mus musculus; ChIP-Seq - SRA - NCBI [Internet]. [cited 2022 Sep 13]. Available from: https://identifiers.org/insdc.sra:SRR5085154

92. GSM2418859: WT_Rad21_ChIPSeq; Mus musculus; ChIP-Seq - SRA - NCBI [Internet]. [cited 2022 Sep 13]. Available from: https://identifiers.org/insdc.sra:SRR5085155

93. GSM2418860: WT_CTCF_ChIPSeq; Mus musculus; ChIP-Seq - SRA - NCBI [Internet]. [cited 2022 Sep 13]. Available from: https://identifiers.org/insdc.sra:SRR5085156

94. GSM2418860: WT_CTCF_ChIPSeq; Mus musculus; ChIP-Seq - SRA - NCBI [Internet]. [cited 2022 Sep 13]. Available from: https://identifiers.org/insdc.sra:SRR5085157

95. Reiff SB, Schroeder AJ, Kirli K, Cosolo A, Bakker C, Mercado L, et al. The 4D Nucleome Data Portal: a resource for searching and visualizing curated nucleomics data [Internet]. bioRxiv. 2021 [cited 2022 Apr 11]. p. 2021.10.14.464435. Available from: https://www.biorxiv.org/content/10.1101/2021.10.14.464435v1

