## Additional File 1 for "MoDLE: High-performance stochastic modeling of DNA loop extrusion interactions"

#### Supplementary text

##### 1. MoDLE: design overview

One of the motivations for developing MoDLE is the need for a high-performance in-silico model of DNA loop extrusion. In this day and age writing high-performance software almost always requires developers to target parallel and heterogeneous hardware architectures, as a single-threaded, sequential application can only take advantage of a miniscule fraction of the performance offered by modern hardware. Given this, it was clear from the beginning that in order to meet our performance target we would need to design MoDLE so that it could efficiently run on machines with many CPU cores, ideally splitting the computation in many small independent work units that can be executed on different CPU cores requiring little or no synchronization.
The version of MoDLE presented in this manuscript uses a producer-consumer architecture, where a single producer communicates with multiple consumers through asynchronous message passing.
In this programming model we have a single thread with the role of producer, here and onwards referred to as the main thread. The main thread is responsible for reading input files and generating a set of simulation tasks to be consumed by a pool of worker threads. Tasks are implemented as light-weight C++ structs, and are extremely cheap to generate and consume. A single task contains all the information needed by a worker thread to run a simulation instance, that is simulating DNA loop extrusion on a single chromosome in a specific cell. Simulation instances are for the most part independent from each other and can thus run concurrently. This means that when simulating loop extrusion on the human genome with default settings using 512 cells, MoDLE could in theory run all 11776 simulation instances in parallel. This level of parallelism is orders of magnitude larger than what is available on modern high-performance servers and workstations. This means that MoDLE can efficiently take advantage of all CPU cores available on the host machine, and this is likely to remain true for the foreseeable future.
Another aspect we had to consider during MoDLE's design was memory usage and layout. Modern CPUs take advantage of several techniques such as instruction pipelining, out-of-order and speculative execution to greatly improve the instruction throughput of a single CPU core. Because of this, the bottleneck of many applications running on modern CPUs is rarely instruction throughput, and is instead memory bandwidth. The most important aspect to consider when optimizing a memory-bound application is memory locality of reference, so that the CPU prefetcher can efficiently predict and prefetch data the application will access next, making efficient use of the CPU cache and hiding memory access latencies.
With this in mind we identified a data topology that would minimize the memory used by a

single simulation instance without significantly compromising locality of reference (see Section 3).
Data stored by MoDLE in system memory can be partitioned into two logical classes: data shared across multiple worker threads and data that is owned and accessed by a single worker thread.
A small fraction of shared data is immutable, and can thus be accessed concurrently from different threads without requiring any synchronization. Examples of shared immutable data are the list of chromosomes that comprise the genome that is being simulated or the position, direction and occupancy of extrusion barriers mapping on a given chromosome (see Section 6).
Extrusion barrier state (i.e. occupied or not occupied), LEF positions and states are instead examples of data that must be stored locally by each worker thread (see Sections 5 and 6).
MoDLE does not explicitly model DNA, as this is not necessary with the current design, and would greatly increase MoDLE's memory footprint. Instead, for each chromosome MoDLE only tracks chromosome name, total length and begin/end position (in case chromosomes subsets are being simulated). Extrusion barriers and LEFs are both stored in contiguous memory. Extrusion barriers are sorted by their genomic coordinates, so visiting them in 5'-3' or 3'-5' direction is trivial and can be done in $O(n)$. Sorting needs to be done only once, as extrusion barrier positions are immutable throughout the entire simulation (see Section 6).
LEFs are also stored in contiguous memory but in unspecified order (see Section 5). At the beginning of every epoch, MoDLE builds two indices such that visiting extrusion units moving in forward and reverse direction sorted by their genomic coordinates becomes trivial and can be done in $O(n)$ (see Section 8).
With the ability of visiting extrusion units sorted by their genomic coordinates, all kinds of collision events can be detected by comparing the distance of adjacent elements (e.g. an extrusion unit moving in forward direction and the first extrusion barrier located downstream of said extrusion unit) with the distance the extrusion unit is set to travel during the epoch that is being simulated (see Section 12). If the proposed move is greater than the distance that separates the two entities, then MoDLE predicts a collision may occur in the current epoch. Whether or not a collision actually takes place is determined by a stochastic process (usually a Bernoulli trial). Collisions are processed one type at a time (e.g. first we process collisions between extrusion units and extrusion barriers, next we process collisions between extrusion units moving in opposite directions and so on). For each extrusion unit MoDLE records whether a collision occurred as well as a reference to the entities involved in the collision event (see Section 7). MoDLE also tracks instances where a collision was avoided due to stochastic events (e.g. because two colliding extrusion units were able to bypass each other). Once all collisions have been detected, the proposed moves are updated to satisfy the constraints introduced by collision events (see Sections 12a and 12e-g). The updated moves are then used to update the simulation state before registering contacts, releasing a fraction of the active LEFs, and moving to the next epoch( see Section 6).

The majority of MoDLE's memory budget is used to track molecular contacts generated by DNA loop extrusion. Molecular contacts are stored using a specialized contact matrix data structure described in Section 2, which was developed with three aims: enabling efficient pixel random access, thread safety and low memory overhead.
Storing a single instance of the contact matrix for each chromosome that is being simulated and sharing it across all worker threads allows MoDLE to minimize the amount of memory required to store the state of a single simulation instance. The size of this state varies from

500 KB or less to 5-10MB depending on the number of LEFs and extrusion barriers involved in the simulation instance (see Section 3). The size of the shared contact matrix depends on chromosome size, width of the region along the diagonal that is being represented as well as bin size. With default settings, the contact matrix for chromosome 1 from the human genome requires approximately 120 MB of memory (see Section 2).

#### 2. MoDLE: contact matrix

The main goal when designing the specialized contact matrix data structure used by MoDLE was to take advantage of the nature of DNA loop extrusion to minimize the memory footprint of the contact matrix as well as making most of the class public API thread-safe.
The main strategy used to limit the memory usage of the contact matrix class is to avoid representing regions in the matrix where loop extrusion is unlikely to generate contacts. With default setting the contact matrix is only representing contacts for a 3 Mbp window around the diagonal of the upper triangle of the contact matrix. When a simulation instance tries to set or increment a value outside of the space represented by the contact matrix, the action is effectively a no-op and will lead to a missed update. The number of missed updates are tracked using an atomic counter and a warning is issued if the fraction of missed updates becomes significant. Attempts to fetch the value of a pixel that lies outside of the 3Mbp window is also a no-op and always returns zero.
Using this strategy, storing contacts for chromosome 1 of the human genome using a bin size of 5000 bp and 32-bit integers requires around 120 MB of memory while storing the entire square matrix would require close to 10 GB.
The public API of the contact matrix class is rendered thread-safe through mutex locking. When instantiating the contact matrix class with a certain shape (i.e. the number of rows and number of columns, where the number of rows represents at most 3 Mbp of DNA with default settings), the class constructor will also allocate a vector of mutexes that will be used to protect subsets of the contact matrix from concurrent access from different threads. Access to a given pixel is protected by a single mutex. Given the pixel corresponding to row $r$ and column $c$, to identify which mutex to acquire prior to accessing the pixels, the coordinates $r$ and $c$ are hashed. The output of the hash function is then divided by the number of mutexes protecting the current contact matrix instance, and the remainder of the division is used to index the mutex protecting the pixel that we are about to access. The exact number of mutexes protecting a contact matrix instance is computed based on the matrix shape and the number of available CPU cores. This number is always rounded to the next power of two so that the index of the mutex protecting a given pixel can be computed using bitwise operations instead of slower modulo operations. The number of mutexes is currently clamped between $2$ and $1000 * t$, where $t$ is the maximum number of CPU cores available on the machine where MoDLE is being executed. As an example, on server A (see Table 1), MoDLE allocates up to 256,000 mutexes for each contact matrix, using roughly 10 MB of memory.

#### 3. MoDLE: Private state of a simulation instance

As soon as the first task is consumed by a given worker thread, the worker thread will allocate and initialize the memory required to store the private state of a single simulation instance. The state object itself consists of a struct with several member variables allocated on the stack, such as a 64-bit integer field used to keep track of the simulation epoch and the state of the pseudo-random number generator used by the current simulation instance (see Section 4). The state object also owns and manages a dozen buffers that are used for various purposes throughout the simulation. These buffers are allocated once, then cleared and resized when starting a new simulation instance after reading a new task. The majority of these buffers have a size that depends on the number of LEFs that are being simulated. Given that most of the time chromosomes are processed from the largest to the smallest, and that in MoDLE the number of LEFs is a function of chromosome size, each worker thread is expected to perform several relatively large allocations when processing the first task, and then progressively shrink the buffers, which is a cheap operation to perform, especially compared to growing buffers. This memory allocation strategy allows MoDLE to greatly reduce memory re-allocations, preventing performance issues due to frequent memory allocations and memory fragmentation.

### 4. MoDLE: PRNG and simulation reproducibility

The PRNG algorithm used by MoDLE to model stochastic events can be customized at compile-time. By default MoDLE will be compiled to use xoshiro256++ [1] as PRNG. Alternatively, the Boost or STL implementation of the Mersenne-Twister algorithm (64-bit) can be used.

Each worker thread owns an instance of the PRNG class. After consuming a new task from the queue, worker threads seed the PRNG using a seed computed by hashing chromosome and cell metadata (namely chromosome name, chromosome size and cell ID) with a user-provided seed (optional, specified through the --seed CLI option).

This seeding strategy not only makes simulation results reproducible across runs on the same machine (assuming simulation parameters are unchanged), but also across machines with a different number of CPU cores. It should be noted that result reproducibility is currently not guaranteed across operating systems and on machines using different standard library implementations. Users relying on result reproducibility are encouraged to use the Docker images available on GHCR.io and DockerHub at https://github.com/paulsengroup/modle/pkgs/container/modle or https://hub.docker.com/r/paulsengroup/modle

### 5. MoDLE: modeling LEFs

Loop extrusion factors are modeled as pairs of extrusion units, with one extrusion unit moving in reverse direction and the other in forward. In addition, each LEF keeps track of the epoch corresponding to the latest re-bind event.

Extrusion units are implemented as C++ structs consisting of a single integer representing their position along a chromosome and that implement most comparison operators to allow comparing LEF positions with that of other objects, such as extrusion barriers. Every simulation epoch MoDLE randomly samples a candidate moving distance for each extrusion unit. Moves are drawn from a normal distribution parameterized using the extrusion speed average and standard deviation. Candidate moves correspond to the maximum distance a given extrusion unit will cover during the current epoch (assuming there are no collision events). Upon activation, LEFs are bound to a random position on the chromosome that is

being simulated in the current simulation instance. Binding positions are sampled from a uniform distribution. At the end of every epoch a subset of LEFs are released from the DNA, so that they can be bound at a new location in the next epoch. Whether or not a specific LEF will be released is determined by a Bernoulli trial whose probability of success is computed based on the average LEF processivity and extrusion speed. This probability of success is also affected by whether a LEF is stalled on both sides and for which reason. As an example, with default settings, given a LEF whose extrusion units are stalled by a pair of extrusion barriers in convergent orientation, the probability of release for this specific LEF is reduced by a factor of 5.
With default settings extrusion units have a low probability of bypassing each other. This probability can be controlled through the CLI and can be set to 0 to forbid extrusion unit bypass.

#### 6. MoDLE: modeling extrusion barriers

The version of MoDLE here presented models extrusion barriers based on the CTCF transcription factor using a two-state Markov process.
The two states of the Markov process represent the occupied and not-occupied states of a CTCF binding site by the CTCF transcription factor.
The state of each extrusion barrier at a given epoch is represented with a bitset and is stored outside of the extrusion barrier class and in the thread-local simulation state, so that extrusion barrier objects can be stored once, and shared across all simulation instances.
Each extrusion barrier consists of a position, blocking direction and two transition probabilities. An extrusion barrier instance thus loosely represents a CTCF binding site on the DNA, while its corresponding entry in the bitset signals whether a CTCF protein is bound to this specific binding site.
The recommended way of setting transition probabilities for extrusion barriers is using the data-driven approach described in the Method section, which relies on CTCF and Cohesin ChIP-seq data. Should this data not be available, users can directly specify the transition probabilities on the CLI through the options --extrusion-barrier-bound-stp and --extrusion-barrier-not-bound-stp. Alternatively, users can specify the --extrusion-barrier-occupancy option which is then used to compute the the occupied-to-occupied transition probability with the following formula:

$$p_{BB} = 1 - \frac{p_{UU} - (\pi_B - p_{UU})}{\pi_B}$$

where $p_{BB}$ is the probability that a CTCF site that is currently bound by a CTCF protein will remain in this state in the current epoch, while $p_{UU}$ is the probability that an unoccupied CTCF binding site will remain unoccupied throughout the epoch that is being simulated. Finally, $\pi_B$ is the CTCF binding site occupancy.

#### 7. MoDLE: tracking collisions

MoDLE defines four different kinds of collision events:

- LEF-extrusion barrier collisions (LB) - any collision occurring between a LEF extrusion unit and an extrusion barrier in the bound state.

- Primary LEF-LEF collisions (LL1) - collisions between two extrusion units belonging to two different LEFs and moving towards each other.
- Secondary LEF-LEF collisions (LL2) - collisions between two extrusion units belonging to two different LEFs that are moving in the same direction.
- Collisions between LEFs and chromosomal boundaries (CB).

MoDLE models chromosomal boundaries as impenetrable barriers to prevent LEFs from falling off chromosome ends. Collision events are stored in two vectors, one for extrusion units moving in reverse direction and the other for extrusion units moving forward.
Collision events are represented using unsigned 64-bit integers. Each integer encodes the cause of collision as well as a reference to the entity that caused the collision.
The high bits of a collision event are used to encode the collision type while the lower bits are used to encode the reference to the entity causing the collision. Usually, the reference is an index pointing into one of the vectors of extrusion units or barriers.
Using this encoding scheme makes collision encoding and decoding very efficient, as both operations can be implemented using bitwise operations such as bit shifting and masking.

| | CO | CB | LB | LL1 | LL2 | Index | | | | |
|---|---|---|---|---|---|---|---|---|---|---|
| bit idx | 63 | 62 | 61 | 60 | 59 | … | 3 | 2 | 1 | 0 |
| bit value | 1 | 0 | 1 | 0 | 0 | 0 | 1 | 1 | 0 | 1 |

Legend:
CO - Collision Occurred (1 when collision occurred, 0 when collisions did not occur).
CB - Collision with Chromosomal Boundary.
LB - LEF extrusion barrier collision.
LL1 - LEF-LEF primary collision.
LL2 - LEF-LEF secondary collision.

The above example encodes a collision between the extrusion unit at index $i$ and the extrusion barrier at index 13 (1101 in binary), where $i$ is the index of the collision event in the collision vector.

#### 8. MoDLE: indexing LEFs

MoDLE stores LEFs in contiguous memory in an unspecified order. Extrusion units moving in the same direction are indexed so that they can be visited in 5’-3’ or 3’-5’ direction.
Indexing is accomplished by indirectly sorting extrusion units by their positions and dealing with ties using the LEF binding epoch. Ties are rare events which can occur when a newly bound LEF is assigned a binding position that is already occupied by another extrusion unit.
Indexing is potentially an expensive operation due to the sorting step, as even state of the art general-purpose sorting algorithms run in $O(n \log n)$ where $n$ is the number of LEFs in this case. MoDLE takes advantage of the fact that after the first epoch only a few index entries need to be updated to significantly reduce the time complexity of the sorting step. In other words, traversing extrusion units using the index from the previous epoch yields a sequence of extrusion units that is for the most part already in the correct order. Extrusion units are indirectly sorted using Splitsort [2], an adaptive sorting algorithm that can take

advantage of the level of pre-sortedness of extrusion units indexed using the index from the previous iteration. In brief, given a collection of items to be sorted, Splitsort removes a number of items from the collection, splitting the collection in two partitions: a partition containing already sorted items, and a partition with items yet to be sorted. The unsorted partition is then sorted using a traditional sorting algorithm (Quicksort in the Splitsort implementation used by MoDLE). Finally the two sorted partitions are merged back into the original sequence. Splitsort is not a stable sorting algorithm, meaning that the order of ties in the sorted collection is not deterministic. MoDLE uses LEF activation epoch to make Splitsort practically stable. Let us consider the following scenario: two LEFs $l_1$ and $l_2$ have been assigned to the same chromosome in the same simulation instance. At the end of epoch $e_0$, $l_1$ is active and extruding: its forward-moving extrusion unit $ef_1$ is located at position $p$. At this time, $l_2$ is inactive (i.e. not bound to the chromosome). During the next epoch ($e_1$), $l_2$ becomes active and is bound to position $p$, the same position as $ef_1$. In this scenario $ef_1$ and $ef_2$, the forward-moving units of $l_1$ and $l_2$ respectively are tied. The tie is resolved by observing that $ef_2$ became active after $ef_1$ reached position $p$, and that thus reached position $p$ after $ef_1$. Because of this, $ef_2$ is ranked lower than $ef_1$. The case where two extrusion units become active at the same time and bind to the same position is not explicitly handled, as under normal circumstances these are extremely rare events.

#### 9. MoDLE: generating molecular contacts

At the end of each epoch after the burn-in phase, MoDLE selects a number of LEFs by random sampling with replacement for contact registration. The genomic coordinates of these LEFs will be used to generate molecular contacts. Contact sampling rate can be tweaked through the --contact-sampling-interval parameter, which sets the average number of base pairs a LEF extrudes between two subsequent sampling events (assuming loop extrusion proceeds unencumbered). The default value for this parameter is 50 kbp.
MoDLE can sample two kinds of contacts:

- Molecular contact directly mediated by a LEF, that is contacts between the two regions brought together by the extrusion units of a LEF. This type of contact will very often occur in correspondence of stripes and dots, see Fig 2A.
- Contacts between regions within the same loop. This type of contact almost always occurs inside squares on the diagonal corresponding to TADs, see Fig 2B.

Furthermore, contacts can be randomized by adding noise sampled from a GEV distribution whose location, scale and shape parameters can be customized from the CLI.
At the time of writing, MoDLE supports the following contact sampling strategies:

- Loop only.
- TADonly.
- TAD + loop.
- Loop only with noise.
- TAD only with noise.
- TAD + loop with noise.

With default settings, MoDLE will, on average, sample 5 TAD contacts for every loop contact. This ratio was determined empirically, with the aim of generating contact matrices resembling that produced by Hi-C and Micro-C. The ratio between the two types of contacts

can be set through the --tad-to-loop-contact-ratio CLI option.
Let us consider LEF $l$ with extrusion units $e_1$ and $e_2$, located ad positions $p_1$ and $p_2$ respectively, where $p_1 \leq p_2$.
To sample one loop contact for LEF $l$, random noise from GEV is added to positions $p_1$ and $p_2$ producing positions $p_1^*$ and $p_2^*$ respectively. In case contact randomization was disabled by the user, a noise of 0 is added. Next, positions $p_1^*$ and $p_2^*$ are mapped to their respective bins in the contact matrix $b_1$ and $b_2$ by dividing them by the bin size $bs$. Finally the value of the pixel corresponding to $b_1$ and $b_2$ is increased by 1.
The procedure to sample one TAD contact is identical to the one described above except for the last two steps. After computing $p_1^*$ and $p_2^*$, MoDLE will sample two integral numbers between $p_1^*$ and $p_2^*$ from a uniform distribution, $p_3^*$ and $p_4^*$ respectively. $p_3^*$ and $p_4^*$ are then mapped to their corresponding bins $b_3$ and $b_4$, and the value of the corresponding pixel is increased by one.
In MoDLE v1.0.0-rc.7 the CLI option --track-1d-lef-position can be used to track LEF coordinates in 1D space. This information is written to disk in BigWig format. LEF occupancy is recorded similarly as for loop contacts, except that the two positions are mapped to their respective bins, and counts in a 1D vector corresponding to these two bins are incremented accordingly.

#### 10. MoDLE: generating candidate moves

At the beginning of every epoch each active extrusion unit is assigned a candidate move. Moves are randomly sampled from a normal distribution with $\mu = \bar{s}$ and $\sigma = 0.05\mu$, where $\bar{s}$ corresponds to the average extrusion speed. All three parameters $\mu$, $\sigma$ and $\bar{s}$ can be tweaked through MoDLE's CLI. This strategy can lead to unlikely moves for consecutive extrusion units, which is why once all moves have been sampled, moves for consecutive extrusion units are adjusted as follows.
Let us consider two consecutive extrusion units $ef_1$ and $ef_2$ both moving in forward direction. Extrusion units are currently located at positions $p_1$ and $p_2$ respectively, and at a distance $d_{12} = p_2 - p_1$ with $p_1 < p_2$ . Let us also consider two candidate moves $m_1$ and $m_2$ which have been assigned to $ef_1$ and $ef_2$ respectively.Applying these moves to their respective units will bring $ef_1$ and $ef_2$ to positions $p_1^* = p_1 + m_1$ and $p_2^* = p_2 + m_2$ respectively. Finally let us stipulate that $p_1^* \geq p_2^*$ (i.e. $ef_1$ will surpass $ef_2$ by the end of the current epoch), and that $ef_2$ is the only entity located between positions $p_1$ and $p_1^*$ (i.e. $ef_1$ cannot be stalled by an extrusion barrier). The scenario described above does not seem very plausible. A more plausible scenario is that after approaching $ef_2$, $ef_1$ pushes against $ef_2$, leading to a transient increase of the extrusion speed of $ef_2$, lasting until the end of the current epoch. To improve simulation realism, MoDLE, candidate moves are adjusted to reflect the second

scenario: $m_2$ is adjusted to $m_2 = p_1^* - (p_2 + 1)$, so that at the end of the current epoch $ef_2$ will be located immediately downstream of $ef_1$. A similar procedure is used to adjust the moves of adjacent extrusion units moving in reverse direction. The above procedure is used to adjust moves for all pairs of consecutive extrusion units moving in the same direction. Finally, moves are clamped such that no extrusion unit will move past chromosomal boundaries.

#### 11. MoDLE: burn-in phase

With the intent of progressively activate LEFs, as well as limiting the chance of having batches of LEFs in highly correlated states (e.g. groups of LEFs all being bound or released around the same epoch), MoDLE starts new simulation instances by running a burn-in phase.

The burn-in phase consists of two stages: a first stage where LEFs are activated and bound to chromosomes for the first time, and a second stage where all LEFs are active and extruding. During both stages molecular contacts are not recorded. Thus LEF states and positions during burn-in are not directly observable in the output contact matrix.

The burn-in phase is implemented as a bit of extra logic running at the beginning of every epoch. The duration of the first burn-in stage is controlled through the --burnin-target-epochs-for-lef-activation option, which controls the number of epochs over which LEFs are activated. By default --burnin-target-epochs-for-lef-activation is computed from the average LEF processivity and extrusion speed such that by the time the last LEF is activated, the first LEF to have been activated has been released 5 times (approximately 190 epochs with default settings).

Let us consider a chromosome $c$ of size $s_c = 10\ Mbp$ with a LEF density $\lambda_{LEF} = 10\ LEFs\ /\ Mbp$. Let us also assume MoDLE was configured to activate LEFs during a period of 25 epochs, $\Delta A_{LEF} = 25$. In this scenario, during the first burn-in stage, MoDLE will activate on average 4 LEFs every epoch. The exact number of LEFs to be activated in a given epoch are sampled from a Poisson distribution with $\mu = \frac{\lambda_{LEF} s_c}{\Delta A_{LEF}}$. LEFs are activated by drawing a genomic coordinate on chromosome $c$ from a uniform distribution, and binding both LEF extrusion units at that position (see Section 5).

The second burn-in stage begins as soon as all LEFs have been activated and have extruded some amount of DNA at least once. The purpose of the second burn-in phase is to let the state of LEFs that have been activated during the same epoch to de-correlate from one another, as well as allowing the formation of some large loops of DNA. MoDLE will execute the second burn-in phase for a variable number of epochs, until the average loop size has stabilized. Users can set an upper bound to the duration of the second burn-in phase through the CLI.

The stages through which MoDLE steps every simulation epoch are the same, regardless of whether or not the burnin phase has been completed. The only difference is that steps involved in contact registration are not executed during the burn-in phase.

### 12. MoDLE: automatic parameter adjustment

MoDLE v1.0.0-rc.7 introduces functionality to automatically adjust certain parameters when changing the average extrusion speed directly (i.e. through --rev-extrusion-speed or --fwd-extrusion-speed CLI options) or indirectly (e.g. by changing resolution, as the overall average extrusion speed is set to 1.6 times the resolution by default).
Automatic parameter adjustment is enabled by default, but can be disabled through the --no-normalize-probabilities CLI option.
As of MoDLE v1.0.0-rc.7, the following parameters are adjusted:

- --extrusion-barrier-bound-stp
- --extrusion-barrier-not-bound-stp
- --probability-of-lef-bypass
- --lef-bar-major-collision-prob
- --lef-bar-minor-collision-prob

To motivate the existence of this functionality, let us consider the following scenario.
A MoDLE simulation is started with the following CLI options:

- --extrusion-barrier-occupancy=0.5 ($\pi_B$)
- --extrusion-barrier-not-bound-stp=0.5 ($P_{UU}$)
- --rev-extrusion-speed=4kbp ($s_{REV}$)
- --fwd-extrusion-speed=4kbp ($s_{FWD}$)
- --probability-of-lef-bypass=0.1 ($P_{bypass}$)

In this scenario, active LEFs are expected to extrude an average of 8kbp of DNA per simulation epoch (assuming no collisions take place). With the above extrusion barrier transition probabilities, we can expect barriers to switch between the occupied and not-occupied states once every epoch (on average). Assuming LEFs being simulated are Cohesin-like (extrusion speed=2.1kbp/s [3]), this means extrusion barriers switch states once every 3.81 seconds. Following the same logic, LEF-LEF collisions are expected to resolve themselves within ten epochs on average, which amount to about 50 seconds.
Let us now consider another simulation scenario, where:

- --extrusion-barrier-occupancy=0.5
- --extrusion-barrier-not-bound-stp=0.5
- --rev-extrusion-speed=500bp
- --fwd-extrusion-speed=500bp
- --probability-of-lef-bypass=0.1

With an overall average extrusion speed of just 1kbp/epoch, extrusion barriers are now changing state once every 0.48 seconds, while LEF-LEF collisions are expected to only last 4.76 seconds on average. The two scenarios are clearly not equivalent, in fact simulating at lower extrusion speeds will lead to weaker stripes (especially those produced by already weak barriers).
The automated parameter adjustment here described addresses the above issue.
We introduce a normalization factor $n$, which is equal to twice the default extrusion speed ($n = 8000$). The factor $n$ expresses that transition and collision probabilities specified through the CLI are expressed assuming unencumbered LEFs extrude an average of $n$ base pairs of DNA every epoch. The actual transition and collision probabilities used by MoDLE are then recomputed based on the normalization factor.
We can now revisit the two scenarios described earlier.

In the first scenario probabilities require no adjustment, as the average extrusion speed is already 8kbp/epoch.
In the second scenario, probabilities need to be updated, as the average extrusion speed and $n$ are different. As an example, to update $P_{bypass}$, first we compute the ratio between the average extrusion speed an the normalization factor $n$: $r = \frac{s_{REV} + s_{FWD}}{n} = \frac{500 + 500}{8000} = 0.125$, the adjusted $P_{bypass}$, $P^*_{bypass}$ is then computed as $P^*_{bypass} = r \cdot P_{bypass}$, which is the probability of observing $n$ independent events with equal probabilities.
For the extrusion barrier transition probabilities, first we compute $P_{BB}$ from $\pi_B$ and $P_{UU}$, then we compute the adjusted $P_{BB}$ as $P^*_{BB} = e^{ln(P_{BB})\cdot r}$. Finally we also compute the adjusted $P_{UU}$, $P_{UU}$ from $P^*_{BB}$ and $\pi_B$. Given that extrusion barriers are simulated with a 2 state Markov process, the transition probability adjustment is equivalent to modeling extrusion barriers using a continuous-time Markov chain.

### 13. MoDLE: detecting and processing collisions

#### a. Overview

MoDLE approach to simulate loop extrusion can be summarized as:

- Proposing a set of candidate moves (see Sections 5 and 9).
- Detecting collisions and adjusting the candidate moves to satisfy the constraints introduced by collision events (see Section 12).
- Extrude DNA (i.e. advance extrusion units by their respective adjusted moves).

Collision detection is by far the most complex part of the model. This is one of the reasons why MoDLE does not attempt to detect all kinds of collisions at the same time.
To detect whether a collision event has already occurred or will occur in the current epoch, candidate moves are summed to the positions of forward-moving extrusion units or subtracted to the position of units moving in reverse direction. Then, MoDLE checks whether advancing extrusion units in this way will cause units to collide with other entities (e.g. extrusion barriers or other extrusion units). MoDLE can be parametrized such that certain collision events are not deterministic. For example, with default settings, two extrusion units that are on track to collide with one another during the current epoch have a probability of bypassing each others $p_{bypass} = 0.1$. Whether or not a stochastic collision event will take place in the current epoch is determined by a Bernoulli trial with probability of success directly set or computed from the input parameters. The occurrence and avoidance of collision events is recorded in a collision mask using the approach outlined in Section 7.
For reasons that will become clear in the next paragraphs, the order in which collisions are detected and processed is important. In the implementation here presented collisions are processed in the following order:

- Detect collisions with chromosomal boundaries (i.e. extrusion units that are located at, or are about to reach the 5' or 3'-end of a chromosome).
- Detect collisions between extrusion units and extrusion barriers (LB collisions).
- Detect primary LEF-LEF (LL1) collisions, that is collisions between extrusion units belonging to different LEFs and moving towards each other.
- Correct moves for extrusion units involved in LB collisions.
- Correct moves for extrusion units involved in LL1 collisions.

- Detect secondary LEF-LEF (LL2) collisions and correct moves immediately. LL2 collisions are defined as collisions between extrusion units moving in the same direction and belonging to different LEFs.
- Deal with rare edge-cases that have not been properly handled in the previous steps.

#### b. Detecting extrusion units at chromosomal boundaries

MoDLE models the 5' and 3' ends of chromosomes as impenetrable barriers to prevent LEFs from falling off the simulated chromosome. Thanks to the LEF indexing strategy described in Section 8, detecting collisions with chromosomal boundaries is trivial. First, let us observe that collisions with the 5'-end can only involve extrusion units moving in reverse direction, while collisions with the 3'-end can involve forward-moving units only.
Thus, we can detect all collisions with the 5'-end by traversing reverse-moving extrusion units in 5'-3' order, and looking for extrusion unit $er_i$, which is the last extrusion unit in 5'-3' order that satisfies the following two conditions:

- After extrusion, $er_i$ will be located upstream of unit $ef_0$, which is the first forward-moving extrusion unit in 5'-3' order.
- $er_i$ position after extrusion will be equal to $p_{5P}$, the position of the 5'-end (i.e. position 0 when simulating entire chromosomes).

The first condition is necessary, as if there are one or more extrusion units moving in opposite direction between $er_i$ and the 5'-end, then it is possible that $er_i$ will be stalled by a LL1 collision (see Section 12d). Note that the second condition does not handle the case where $er_i < p_{5P}$ , as this cannot happen (see Section 10).
The same logic can be applied to detect collisions with the 3'-end.Collisions detected in this step are registered in the appropriate collision mask (see Section 7). The index portion of the encoded collision event is set to 5 and 3 for collisions with the 5'-end and 3'-end respectively.

#### c. Detecting LEF-barrier (LB) collisions

To simplify the implementation of this step, MoDLE first detects LB collisions involving reverse-moving extrusion units, and later LB collisions for forward-moving units.
Let us consider the algorithm to detect LB collisions with reverse units. A similar algorithm is used to detect LB collisions involving forward-moving units. Throughout the rest of the section, the term extrusion unit implicitly refers to reverse-moving extrusion units (unless otherwise specified).
The algorithm for LB collision detection for reverse-moving units operates on the following data:

- $E$ - Vector of extrusion units. Units are stored in an unspecified order.
- $I$ - Index vector used to traverse units in $E$ sorted by their genomic coordinates.
- $M$ - Vector of candidate moves.
- $B$ - Vector of extrusion barriers sorted in 5'-3' direction.
- $BS$ - Bitset representing extrusion barrier states (i.e. whether a given barrier is occupied in the current epoch and simulation instance).
- $C$ - Collision mask.

All of the above vectors except $B$ and $B_S$ have size $n$, where $n$ corresponds to the number of LEFs that are being simulated in the current simulation instance. $B$ and $B_S$ are of size $m$, where $m$ is equal to the number of extrusion barriers mapping on the chromosome that is being simulated.
Finally let us consider indices $i$, $j$ and $k$, which point to extrusion barrier $B_i$, index $I_j$ and extrusion unit $E_k$ respectively.
First, indices $i$, $j$ and $k$ initialized such that $i = 0$, $j = 0$ and $k = I_j = I_0$.
Next, we enter a loop that continues until $i = n$ or $j = m$.
Each loop iteration the following steps are performed:

1. If barrier $B_i$ is not occupied (i.e. $BS_i = 0$), jump to step 8.
2. Index $j$ is incremented until $k = I_j$ points to the first extrusion unit located downstream of $B_i$.
3. Compute distance $d_{E_k B_i}$ between unit $E_k$ and barrier $B_i$ as $d_{E_k B_i} = E_k - B_i$ .
4. Distance $d_{E_k B_i}$ is compared with move $M_k$, which is the candidate move for extrusion unit $E_k$.
5. If $M_k < d_{E_k B_i}$, then $E_k$ will not collide with $B_i$ in the current epoch and no further processing of this pair of extrusion barrier and unit is needed. Continue from step 8.
6. If $M_k \geq d_{E_k B_i}$, then it is possible that a collision between $E_k$ and $B_i$ will occur during the current epoch. Whether or not a LB collision will occur depends on the outcome of a Bernoulli trial with probability of success $p_S$, where a success indicates a collision will occur and a failure indicates that the collision will be avoided. $p_S$ is equal to $p_{block\ major}$ if the motif underlying $B_i$ points against the direction of extrusion and $p_{block\ minor}$ otherwise. With default settings $p_{block\ major} = 1$ and $p_{block\ minor} = 0$, meaning that the outcome of the Bernoulli trial is deterministic, and only depends on barrier and extrusion directions.
7. Collision occurrence or avoidance is recorded in the collision mask $C$ at entry $C_k$. In this case, the index portion of the encoded collision $C_k$ is set to $j$, so that later simulation stages can inspect the collision mask and determine which extrusion barrier caused the collision.
8. Increment index $i$ by one and continue from step 1.

Notice how indices $i$ and $j$ are never decremented, meaning that extrusion units and barriers are visited at most once, leading to a worst-case performance of $O(n + m)$.
A similar procedure is followed to detect LB collisions involving forward-moving extrusion barriers. In this case extrusion units are processed in 3'-5' order.

#### d. Detecting primary LEF-LEF (LL1) collisions

The algorithm for LL1 collision detection for reverse-moving units operates on the following data:

- $ER$ and $EF$ - Vector of reverse and forward-moving extrusion units respectively. Both vectors store units in an unspecified order.
- $IR$ and $IF$ - Index vectors for units from $ER$ and $EF$ respectively.
- $MR$ and $MF$ - Vector of candidate moves for $ER$ and $EF$ respectively.
- $B$ - Vector of extrusion barriers sorted in 5'-3' direction.
- $CR$ and $CF$ - Collision masks for $ER$ and $EF$ respectively.

All of the above vectors except $B$ have size $n$, where $n$ corresponds to the number of LEFs that are being simulated in the current simulation instance. $B$ is instead of size $m$, where $m$ is equal to the number of extrusion barriers mapping on the chromosome that is being simulated.
Let us also consider indices $i$ and $j$, which will be used to traverse extrusion units in 5'-3' order using $IR$ and $IF$ respectively.
First, $i$ and $j$ are initialized such that $i = IF_0$ and $j = IR_0$.
Next, we enter a loop that will continue until the end of $ER$ or $EF$ is reached. In the loop, the following steps are performed:

1. Advance index $j$ until $j$ points to the first extrusion unit $ER_j$ that is located downstream of $EF_i$.
2. Advance index $i$ until $i$ points to the last extrusion unit $EF_i$ that is located upstream of $ER_j$.
3. Compute distance $d_{ER_jEF_i}$ between units $ER_j$ and $EF_i$ using $d_{ER_jEF_i} = ER_j - EF_i$.
4. Compare distance $d_{ER_jEF_i}$ with the sum of candidate moves $MR_j$ and $MF_i$.
5. If $MR_j + MF_i < d_{ER_jEF_i}$, then a LL1 collision between $ER_j$ and $EF_i$ during the current epoch is not possible. Continue from step 1.
6. If $MR_j + MF_i \geq d_{ER_jEF_i}$, then a LL1 collision between $ER_j$ and $EF_i$ may take place during the current epoch. This depends on the outcome of a Bernoulli trial with probability of success $p_S = 1 - pb_{LEF}$, where $pb_{LEF}$ is the probability of LEF bypass ($pb_{LEF} = 0.1$ with default settings). If the Bernoulli trial has a negative outcome, then the LL1 collision was avoided. Register collision avoidance in the appropriate collision mask and continue from step 1.
7. The next depends on the current state of units $ER_j$ and $EF_i$, more specifically on whether they can be moved, (i.e. whether one or both of them are already involved in a LB or LL1 collision).
    a. If both units are free to move, register a LL1 collision in the collision masks for entries $CR_j$ and $CF_i$ using $i$ and $j$ for the index part of the encoded collision respectively (i.e. record that $CR_j$ is colliding with $CF_i$ and vice versa). Finally, jump to step 1.
    b. If only one of the colliding extrusion units can be moved (i.e. one of the extrusion units is already involved in another collision), then only register a LL1 collision for the extrusion unit that is free to move.
    In reality this step is more complicated than this, because we need to handle

an edge case where a LB collision needs to be converted to a LL1 collision. This can happen when $ER_j$ and $EF_i$ are located on opposite sides of an extrusion barrier $b$, and one of them is being stalled by the extrusion barrier $b$. Depending on the effective extrusion speed of the two units, the LL1 collision may occur before the LB does. If this is the case, then the LB collision should be replaced with an LL1 collision.

Notice how also in this case, $i$ and $j$ are never decremented, meaning that extrusion units are visited at most once, leading to a worst-case performance of $O(n)$.

#### e. Correcting moves for units involved in LB collisions

Correcting moves to satisfy the constraints introduced by LB collisions is fairly trivial. All we have to do is to loop over the collision masks for reverse and forward moving extrusion units, $MR$ and $MF$ respectively, and look for entries where the appropriate bits are set, namely the CO and LB bits (see Section 7), and adjust the corresponding candidate move.
Let us consider the case where we are adjusting moves for extrusion units moving towards the 5'-end. In this step we operate on the following data:

- $E$ - Vector of extrusion units. Units are stored in an unspecified order.
- $M$ - Vector of candidate moves.
- $B$ - Vector of extrusion barriers sorted in 5'-3' direction.
- $C$ - Collision mask.

Let us stipulate that entry $C_i$ in the collision mask $C$ for $i = 5$ has the following value (see Section 7 for more details on the collision encoding scheme):

| | CO | CB | LB | LL1 | LL2 | Index ($j$) |
|---|---|---|---|---|---|---|
| bit idx | 63 | 62 | 61 | 60 | 59 | 58-0 |
| bit value | 1 | 0 | 1 | 0 | 0 | 15 (decimal) |

This entry encodes a LB collision between the reverse unit $E_i$ and extrusion barrier $B_j$.
With this information, we can now look up the positions for $E_i$ and $B_j$, and compute the corrected move $M_i^*$ as $M_i^* = E_i - B_j - 1$. Advancing unit $E_i$ by $M_i^*$ will thus cause $E_i$ to be located 1 bp upstream of barrier $B_j$.
The same process is repeated for the remaining reverse-moving extrusion units as well as for the forward-moving extrusion units.

#### f. Correcting moves for units involved in LL1 collisions

The procedure to correct moves for extrusion units involved in LL1 collisions is similar to the one described in Section 12e, with two important differences. The first difference is that we are now interested in collision mask entries where both the CO and LL1 bits are set. Second, computing the corrected moves for a pair of colliding extrusion units is a bit more involved.

Let us consider two extrusion units $e_R$ and $e_F$ that are involved in the same LL1 collision event. Unit $e_R$ is located at position $p_R = 500\ bp$ and is moving towards the 5'-end, while unit $e_F$ is located at position $p_F = 150\ bp$ and is moving towards the 3'-end. Let us also stipulate that the candidate moves for $e_R$ and $e_F$ are $m_R = 250\ bp$ p and $m_F = 200\ bp$ respectively.

The position of collision is then calculated with $p_C = p_F + \left(m_F \frac{m_R - m_F}{m_R + m_F}\right)$, and the corrected moves $m_R^*$ and $m_F^*$ are computed as:

$$m_R^* = \begin{cases} p_C + 1 & \text{if } p_C = p_F \\ p_C & \text{otherwise} \end{cases}$$

$$m_F^* = \begin{cases} p_C & \text{if } p_C = p_F \\ p_C - 1 & \text{otherwise} \end{cases}$$

#### g. Processing secondary LEF-LEF (LL2) collisions

In contrast to previous collision detection steps, candidate moves of extrusion units involved in LL2 collisions are corrected immediately after they are detected.
First we process LL2 collisions for reverse-moving extrusion units, and later those involving forward-moving units.
Unless otherwise specified, data and operations outlined in this section are assumed to refer to extrusion units moving in reverse direction.
This step operates on the following data:

- $E$ - Vector of extrusion units. Units are stored in an unspecified order.
- $I$ - Index vector.
- $M$ - Vector of candidate moves.
- $C$ - Collision mask.

In brief, to detect LL2 collisions we loop over consecutive extrusion units in 5'-3' order, and we detect instances where the downstream unit is set to bypass the upstream unit. Given how candidate moves are generated (see Section 10), this can only occur if the upstream unit is being stalled.
Let us consider indices $i = 1$, $j = I_{i-1}$ and $k = I_i$, which will be used to index extrusion units $E_j$ and $E_k$. Let us also consider the candidate moves and entries in the collision mask for $E_j$ and $E_k$, $M_j$, $M_k$ and $C_j$, $C_k$ respectively.
Next, we enter a loop that continues until $i = n$, where $n$ is the number of LEFs in the current simulation instance.
Each loop iteration, the following steps are performed:

1. Keep advancing $i$ until index $j$ points to a stalled extrusion unit and $k$ points to an extrusion unit that is free to move.
2. Compute the position after extrusion for units $E_j$ and $E_k$: $p_j^* = p_j - M_j$ and $p_k^* = p_k - M_k$, where $p_j$ and $p_k$ are the current positions of $E_j$ and $E_k$.

3. If $p_j^* < p_k^*$, then a LL2 collision is not possible. Continue from step 1.
4. $p_j^* \geq p_k^*$, then we a LL2 collision between $E_j$ and $E_k$ may occur during the current epoch. Whether or not this is the case, is determined based on the outcome of a Bernoulli trial with probability of success $p_S = 1 - pb_{LEF}$, where $pb_{LEF}$ is the probability of LEF bypass ($pb_{LEF} = 0.1$ with default settings). If the Bernoulli trial has a negative outcome, then the LL1 collision was avoided. Record collision avoidance in the collision mask and continue from step 1.
5. If the Bernoulli trial from step 4 is successful, then register a LL2 collision with $E_j$ in entry $C_k$ in the collision mask $C$.
6. Lastly, compute the corrected move $M_k^*$ with $M_k^* = p_k - p_j^* - 1$.

A similar procedure is used to detect LL2 collisions for forward-moving extrusion units.

After all LL2 collisions have been processed, we do a final pass over units involved in LL2 collisions to handle an edge-case not handled in the previous step.
Let us consider two consecutive extrusion units moving in forward direction, $ef_1$ and $ef_2$, with $ef_1$ located upstream of $ef_2$. $ef_2$ is stalled due to a LB collision with extrusion barrier $b$. Let us also stipulate that the position of $ef_1$ after move is greater than that of unit $ef_2$ after move, as well as that of barrier $b$: $p_{ef_1}^* > p_{ef_2}^*$ and $p_{ef_1}^* > p_b$. In this scenario, a LL2 collision may have occurred. However, if the LL2 collision was avoided, meaning that the candidate move for $ef_1$ has not been corrected, then extruding $ef_1$ would cause this unit to jump past barrier $b$ (and any other obstacle between $p_{ef_1}$ and $p_{ef_1}^*$ for that matter).

The final pass through units involved in LL2 collisions detect instances like the scenario outlined above, and corrects the move for $ef_1$ such that after extrusion, $ef_1$ will be located 1 bp downstream of $ef_2$.

### References


1. Blackman D, Vigna S. Scrambled Linear Pseudorandom Number Generators. ACM Trans Math Softw. New York, NY, USA: Association for Computing Machinery; 2021;47:1–32.

2. Levcopoulos C, Petersson O. Splitsort—an adaptive sorting algorithm. Inf Process Lett. 1991;39:205–11.

3. Cohesin mediates DNA loop extrusion by a “swing and clamp” mechanism. Cell. Cell Press; 2021;184:5448–64.e22.
