## Additional File 2 for "MoDLE: High-performance stochastic modeling of DNA loop extrusion interactions"

### Supplementary figures

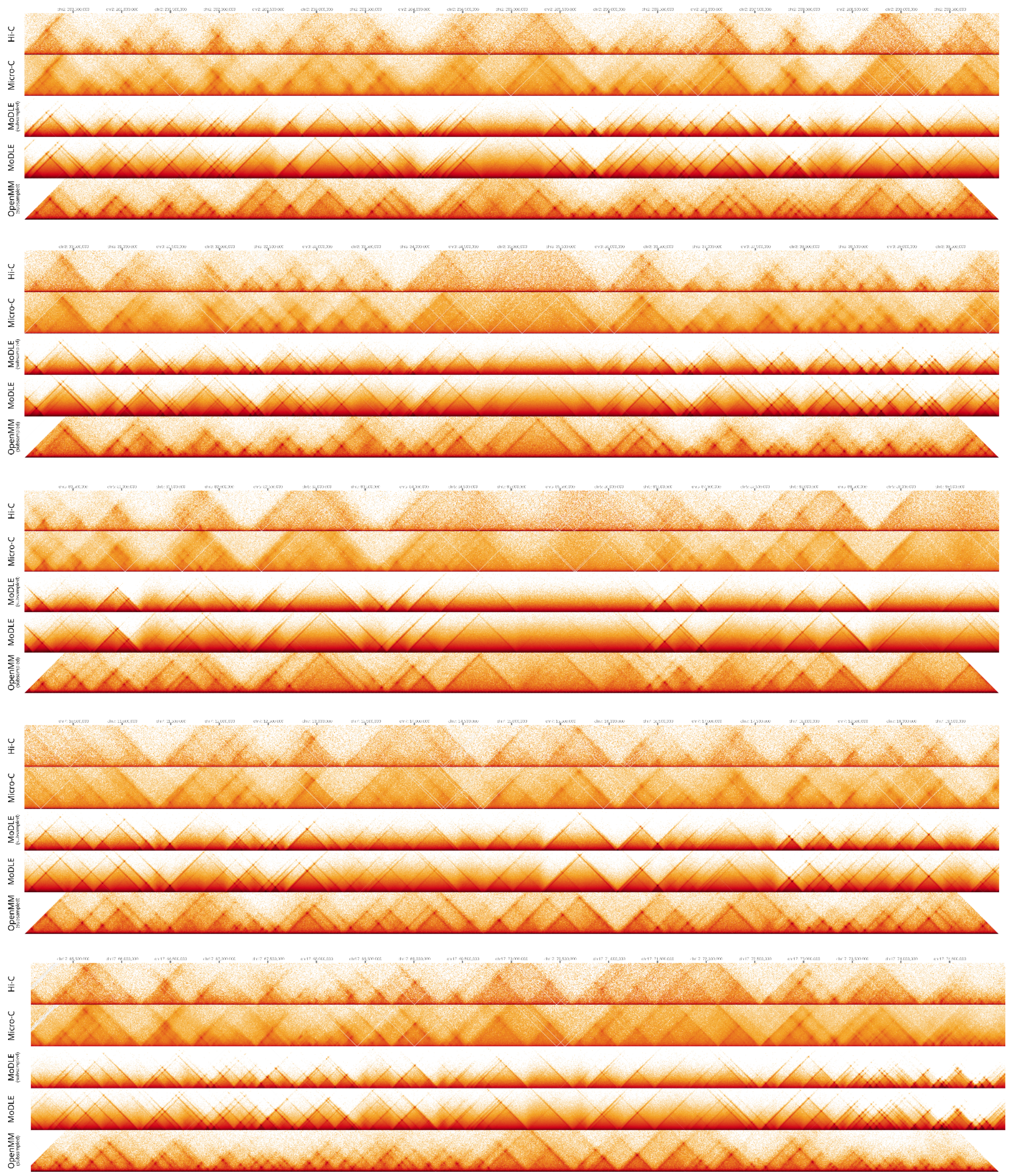


**Supplementary Fig. 1:** Side-by-side comparison of Hi-C and Micro-C data (top two panels),

MoDLE output with and without subsampling (third and fourth panels respectively) and OpenMM output (bottom panels for all selected 10Mbp simulation regions on chromosomes 2, 3, 5, 7 and 17).

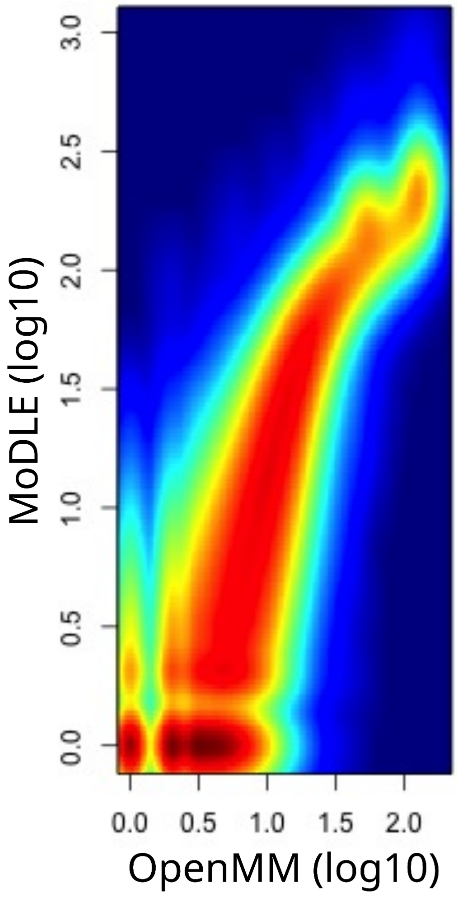


**Supplementary Fig. 2:** Density plot showing frequency (blue = low; red = high) of pixel values in MoDLE (horizontal axis; log10-transformed) vs. OpenMM (vertical axis; log10-transformed). Pearson correlation coefficient is ρ=0.93.

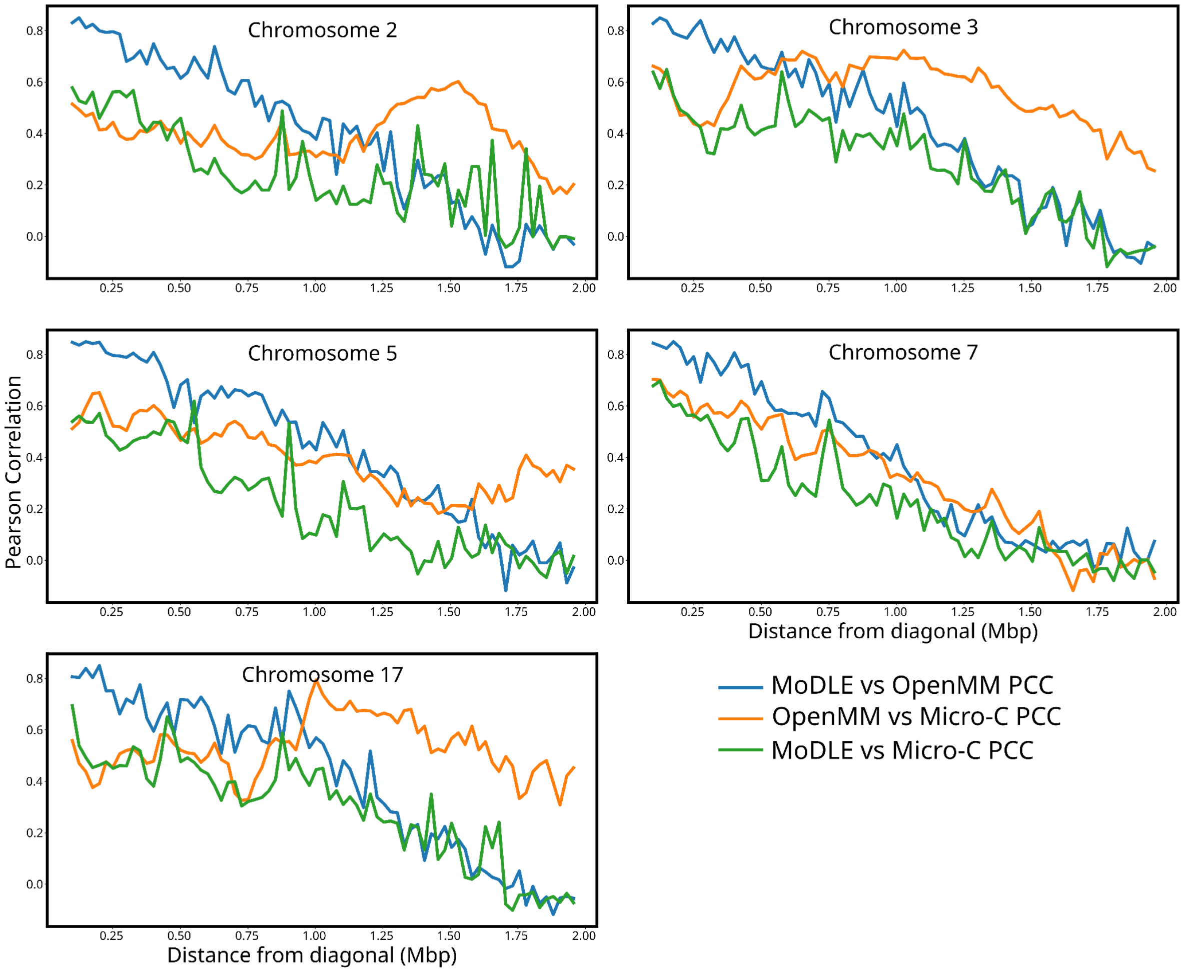


**Supplementary Fig. 3:** Diagonal-by-diagonal correlation plots. Each panel shows the Pearson correlation of MoDLE vs. OpenMM (blue), OpenMM vs. Micro-C (yellow), and MoDLE vs. Micro-C (green) for all selected 10Mbp simulation regions on chromosomes 2, 3, 5, 7 and 17). The correlation is shown for all contact map sub-diagonals from the diagonal to 2Mbp from the diagonal (horizontal axes).

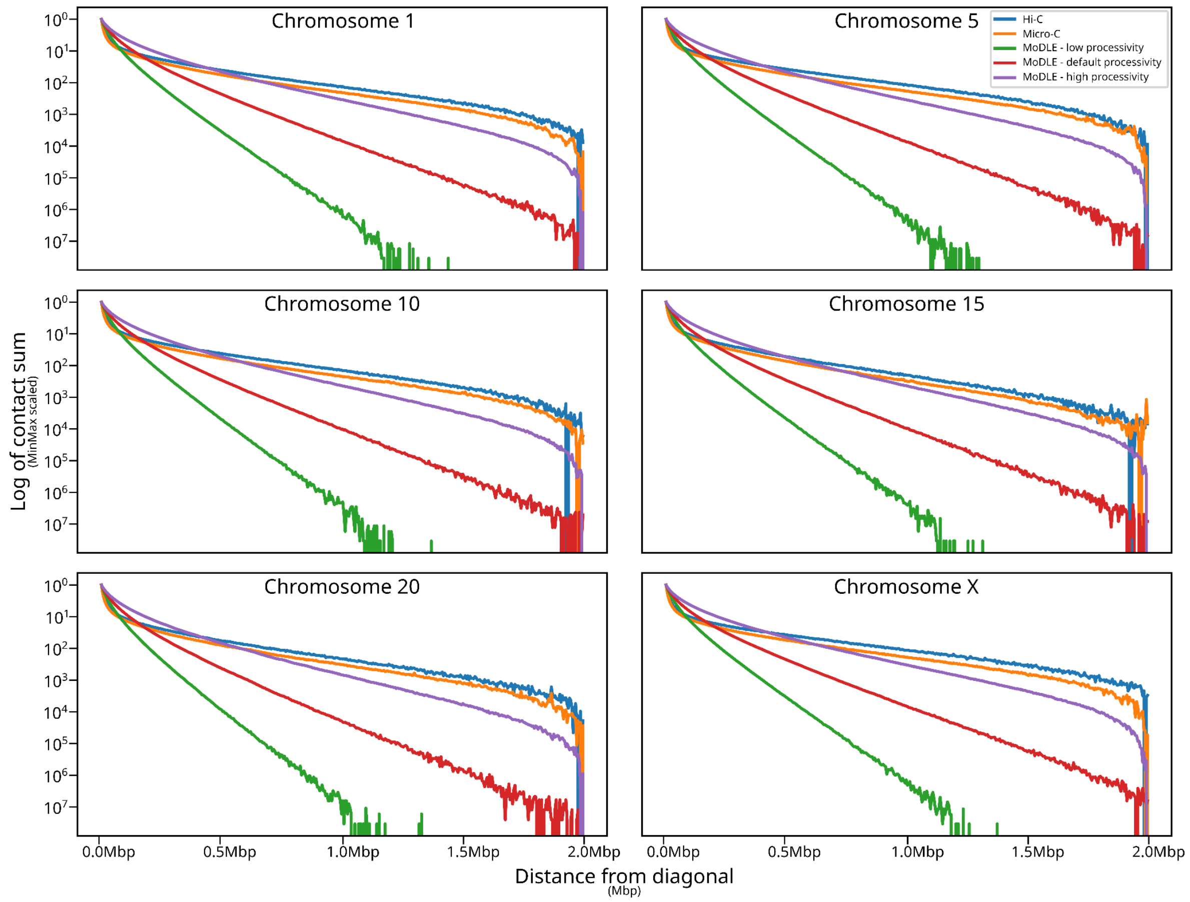


**Supplementary Fig. 4:** Relationship between distance from diagonal (Mbp) and contact frequencies (log of contact sum) for hESC Hi-C data (blue), hESC Micro-C data (yellow), MoDLE using low processivity (150 kbp; green), MoDLE using default processivity (300 kbp; red), and MoDLE using high processivity (600 kbp; purple).

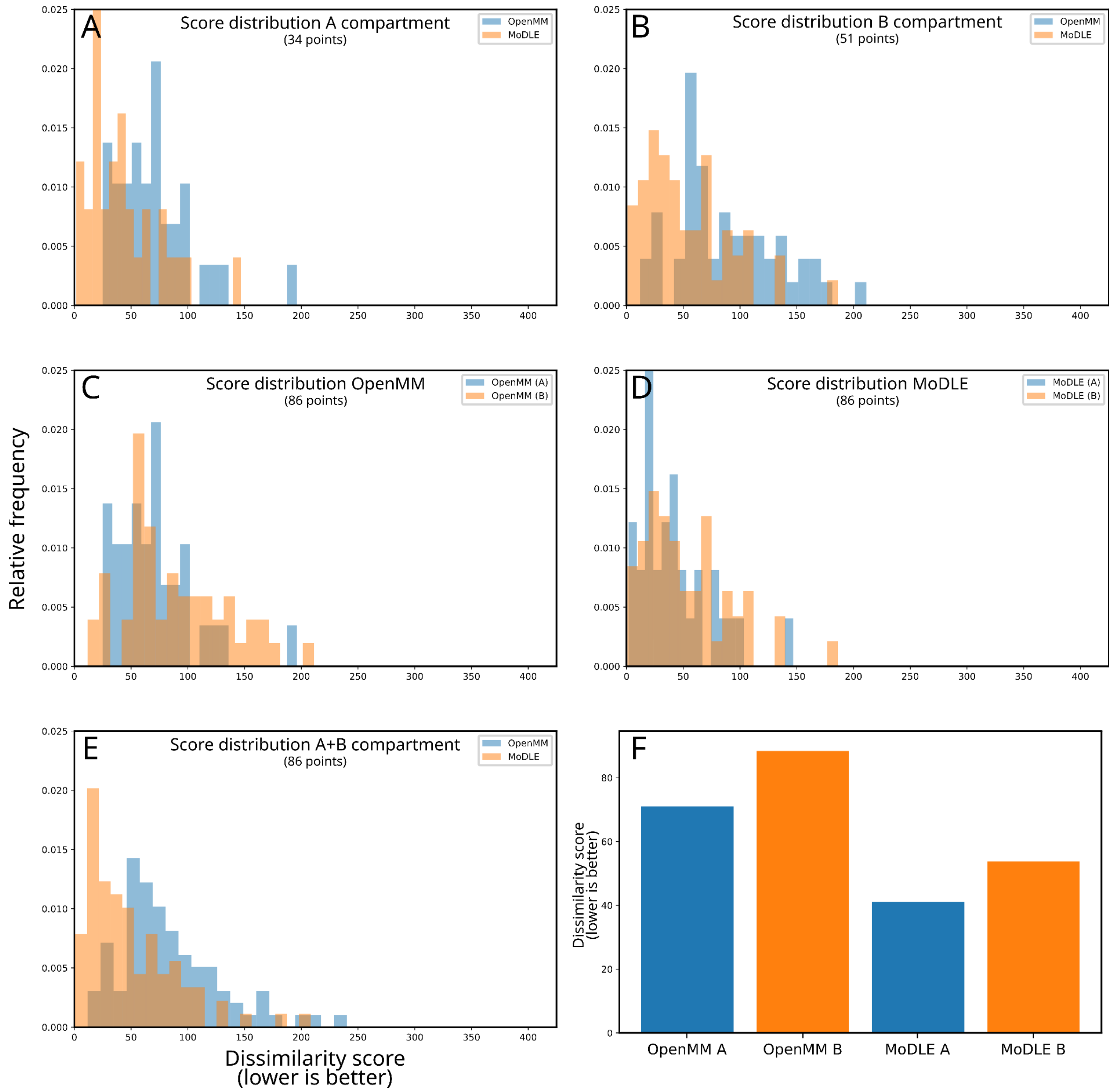


**Supplementary Fig. 5:** Comparison of mean and distribution of dissimilarity scores of MoDLE and OpenMM simulations using H1 hESC Micro-C as reference.
Panel A and B contrast score distribution for MoDLE and OpenMM simulated matrices in A and B compartments respectively.
Panel C and D contrast score distribution of A and B compartments for OpenMM and MoDLE simulated matrices respectively.
Panel E shows the score distribution for OpenMM and MoDLE simulated matrices, regardless of compartment.
Panel F shows the average score for OpenMM and MoDLE simulations in A and B compartments.
Scores are computed as described in Methods (part 6). Before intersecting scores with compartment information, scores are filtered by taking the intersection of the score themselves, the architectural stripes on the reference matrix (stripes annotated with Stripenn) and the genomic regions simulated with OpenMM (chr2:200-210Mbp, chr3:30-40Mbp, chr5:80-90Mbp, chr7:10-20Mbp and chr17:65-75Mbp).

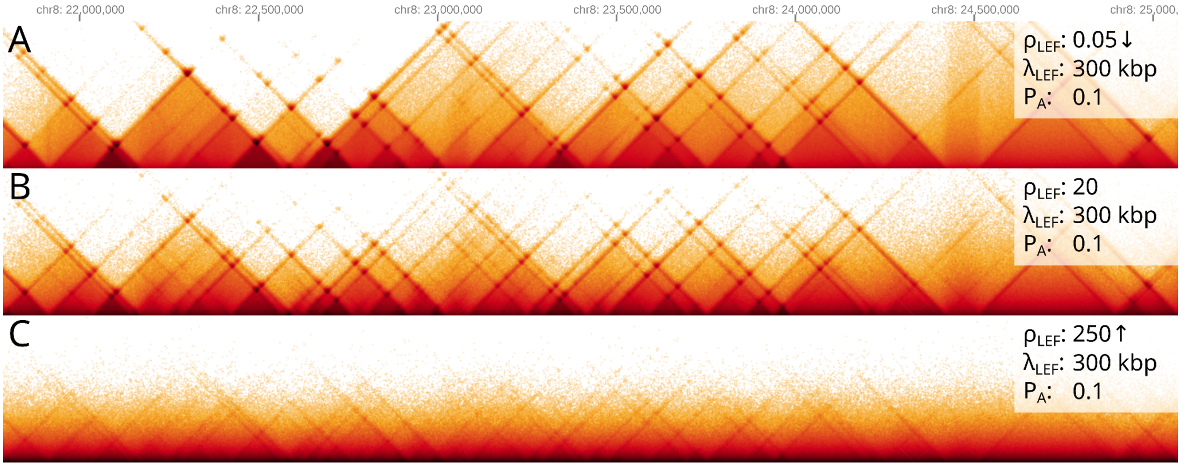


**Supplementary Fig. 6:** Simulated heat maps showing the effect of increasing LEF density ($\rho_{LEF}$).
Higher LEF densities cause contacts to distribute closer to the diagonal. This is due to an increased probability of LEF-LEF collisions. Furthermore, at very high $\rho_{LEF}$ (panel C), stripes and dots appear less defined. This can once again be explained by an increased rate of LEF-LEF collisions, which are stochastic in nature, and do not necessarily occur at or near extrusion barriers.

Legend:
$\rho_{LEF}$: LEF density expressed as the number of LEFs per Mbp of simulated DNA
$\lambda_{LEF}$: Average LEF processivity
$P_A$: Probability of LEF-LEF collision avoidance

↓: Parameter is smaller than the default value
↑: Parameter is larger than the default value

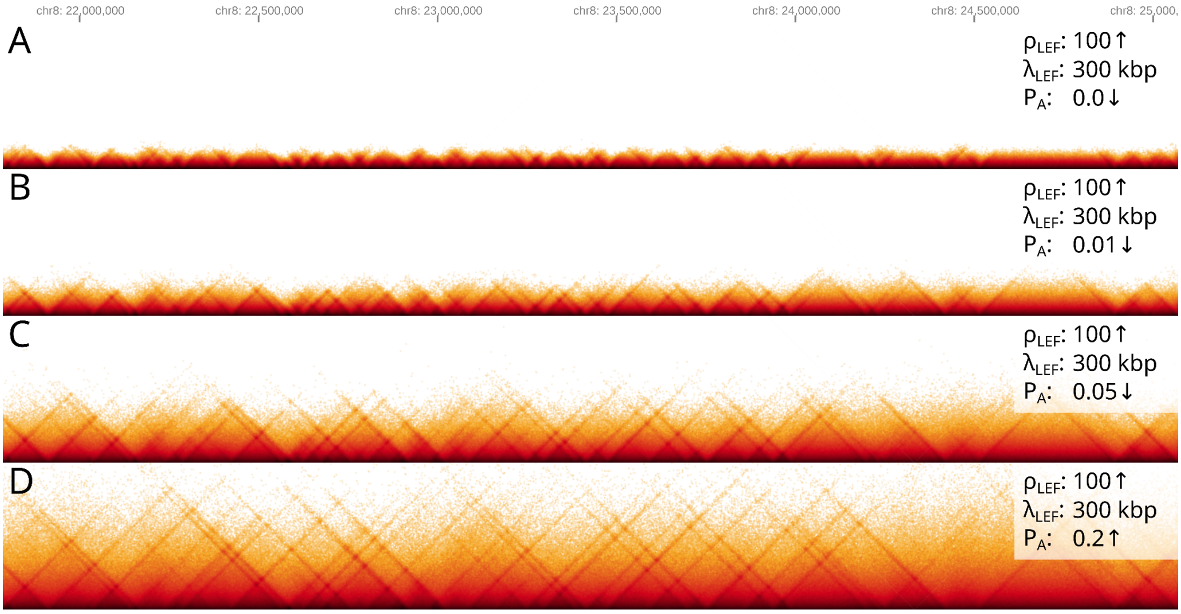


**Supplementary Fig. 7:** Simulated heat maps showing the interplay between LEF density ($\rho_{LEF}$) and probability of LEF-LEF collision avoidance ($P_A$).
This scenario further expands what is shown in Suppl Fig. 4, and shows how increasing $P_A$

can partially counteract very high LEF densities.

Legend:
$\rho_{LEF}$: LEF density expressed as the number of LEFs per Mbp of simulated DNA
$\lambda_{LEF}$: Average LEF processivity
$P_A$: Probability of LEF-LEF collision avoidance

↓: Parameter is smaller than the default value
↑: Parameter is larger than the default value

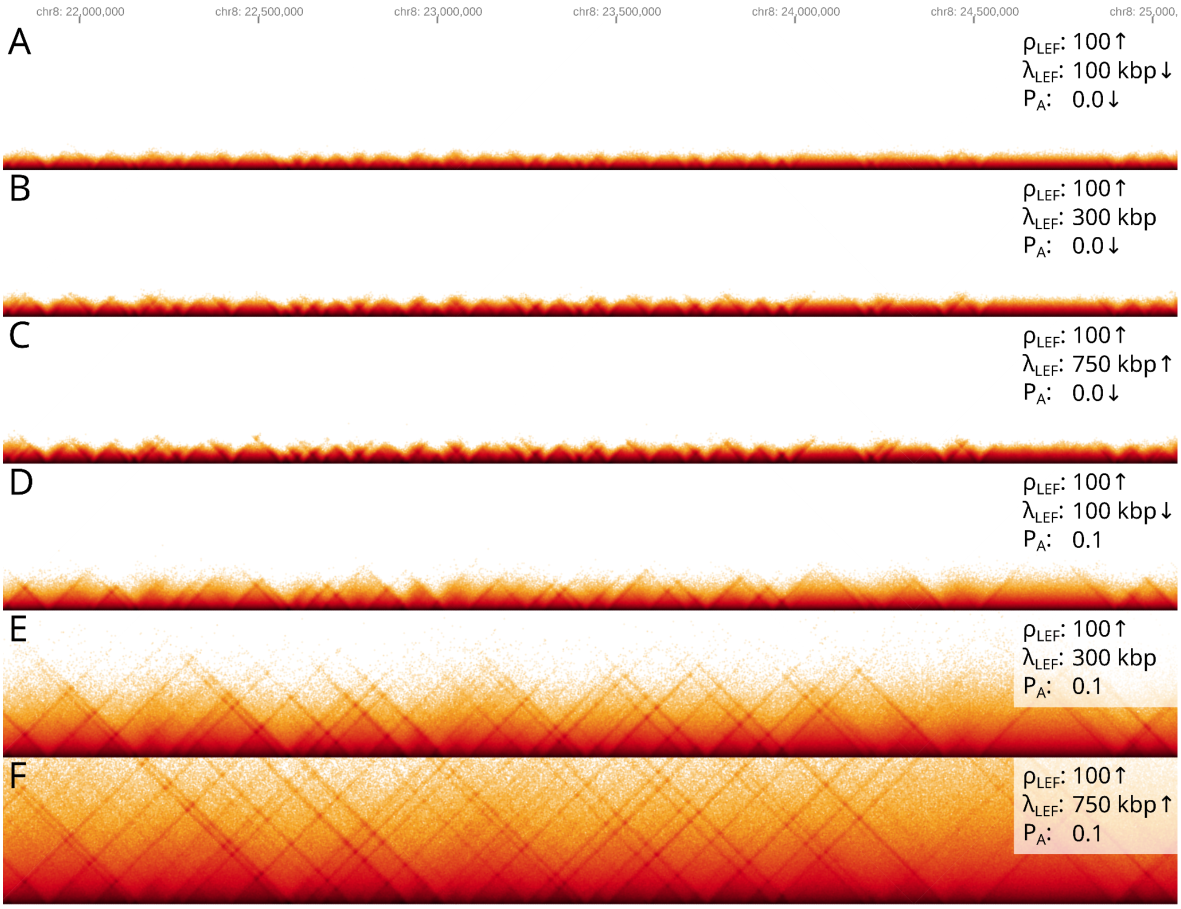


**Supplementary Fig. 8:** Simulated heat maps showing the effect of (dis)allowing LEF-LEF collision avoidance at increasing levels of LEF processivity ($\lambda_{LEF}$).
Increasing $\lambda_{LEF}$ has virtually no effect when the rate of LEF-LEF collisions is high and collisions are deterministic (i.e. LEF-LEF collisions cannot be avoided). Allowing LEFs to bypass one another with a relatively low probability is sufficient to observe progressively longer loops produced by increasingly higher $\lambda_{LEF}$.

Legend:
$\rho_{LEF}$: LEF density expressed as the number of LEFs per Mbp of simulated DNA
$\lambda_{LEF}$: Average LEF processivity
$P_A$: Probability of LEF-LEF collision avoidance

↓: Parameter is smaller than the default value
↑: Parameter is larger than the default value

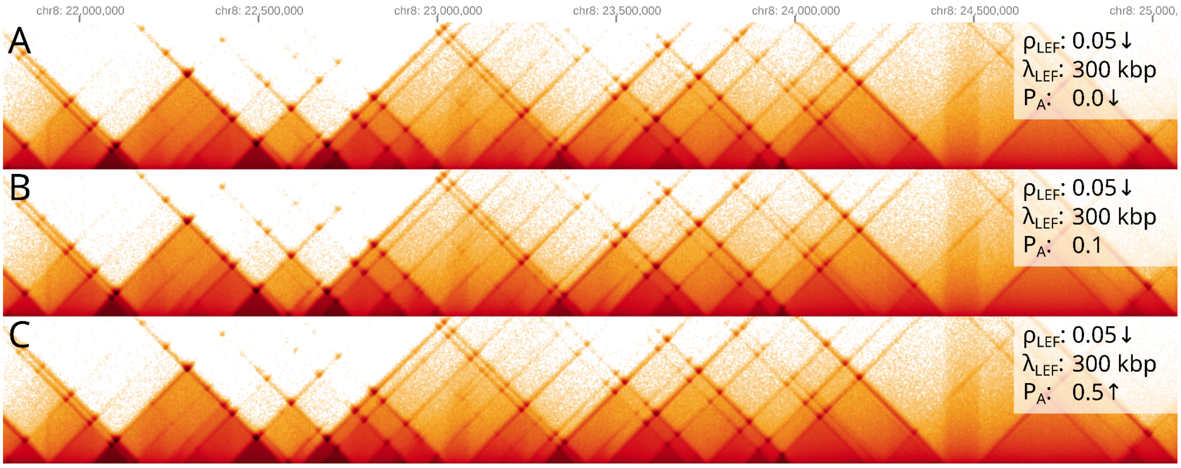


**Supplementary Fig. 9:** Simulated heat maps showing the consequences of simulating with very low LEF densities ($\rho_{LEF}$).
When simulating with very low $\rho_{LEF}$, the probability of two or more LEFs extruding the same region at the same time is approximately 0. Thus, increasing $P_A$ has no effect in practice.
This scenario also highlights one of the consequences of MoDLE simulations not accounting for polymer physics. Very low $\rho_{LEF}$ would likely result in no appreciable contact enrichment due to loop extrusion in vivo or in MD polymer simulations, as once a loop has been extruded, there is plenty of time for the chromatin polymer to relax before the another LEF extrudes through the same region. This is not the case in MoDLE, as loops cease to exist as soon as LEFs are released.

Legend:
$\rho_{LEF}$: LEF density expressed as the number of LEFs per Mbp of simulated DNA
$\lambda_{LEF}$: Average LEF processivity
$P_A$: Probability of LEF-LEF collision avoidance

↓: Parameter is smaller than the default value
↑: Parameter is larger than the default value

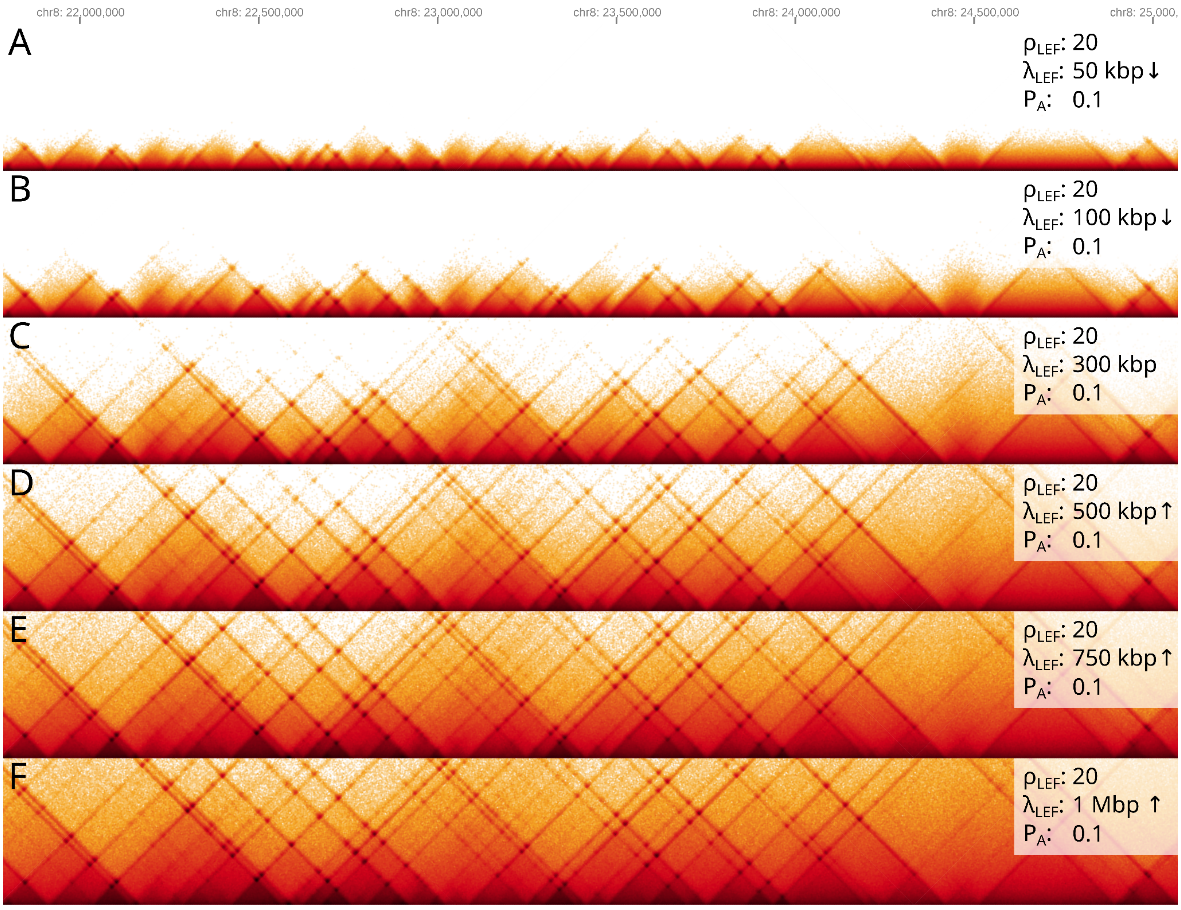


**Supplementary Fig. 10:** Simulated heat maps showing the effect of different LEF processivities ($\lambda_{LEF}$) when LEF density ($\rho_{LEF}$) and probability of LEF-LEF collision avoidance ($P_A$) are left at default.
With realistic LEF concentrations, altering $\lambda_{LEF}$ has the expected effect: higher processivities lead to loops, which result in longer stripes and stronger dots even at relatively high distances from the diagonal.

Legend:
$\rho_{LEF}$: LEF density expressed as the number of LEFs per Mbp of simulated DNA
$\lambda_{LEF}$: Average LEF processivity
$P_A$: Probability of LEF-LEF collision avoidance

↓: Parameter is smaller than the default value
↑: Parameter is larger than the default value

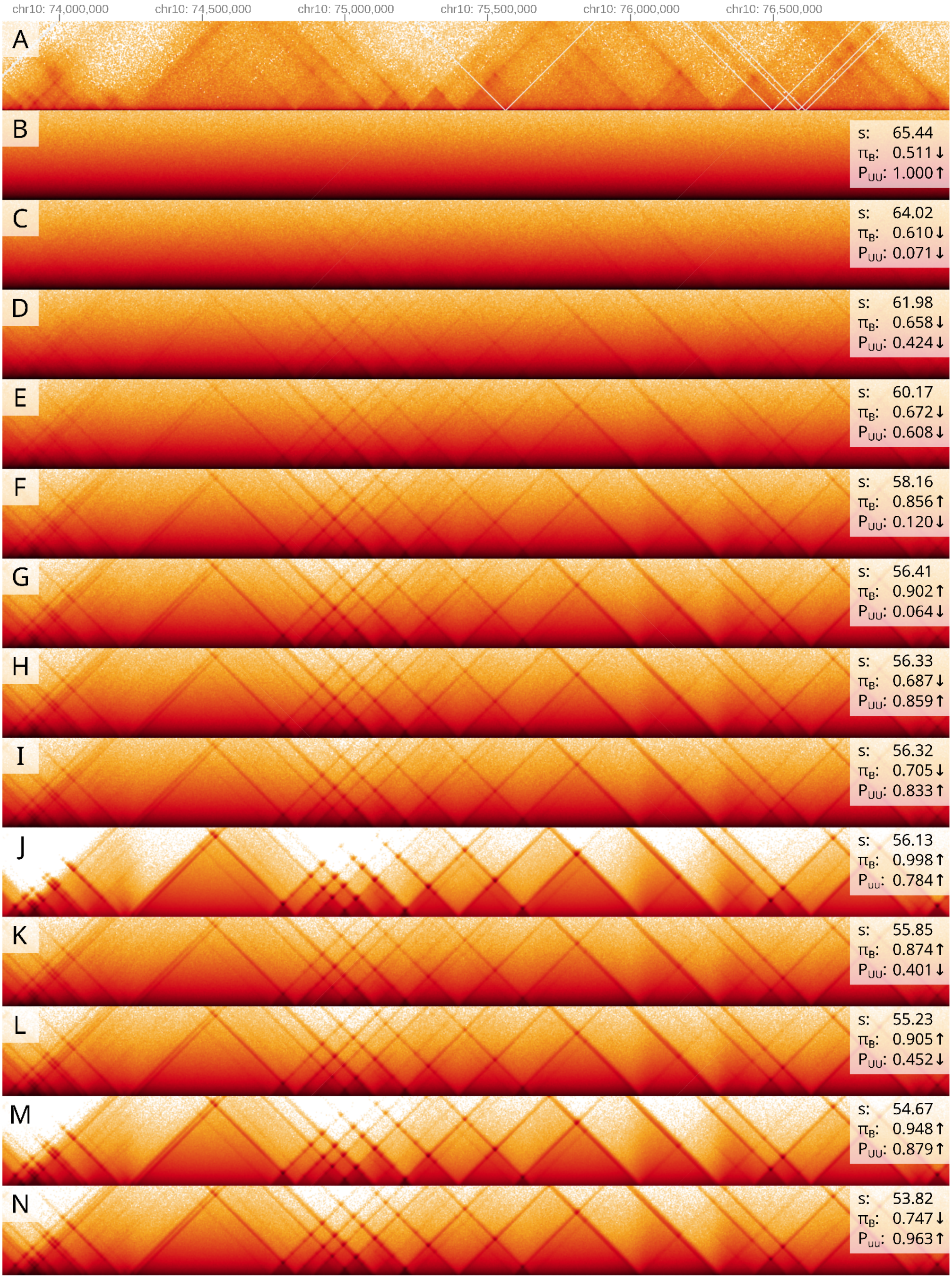


**Supplementary Fig. 11:** Side-by-side comparison of H1-hESC Micro-C data (top panel) and matrices simulated by MoDLE. Panels B-N show a selection of the parameter combinations explored by the Gaussian optimization described in Methods (part 9). On the right, each panel is labelled with its corresponding score $s$, occupancy $\pi_B$ and self-transition probability of the unbound state $P_{UU}$.

Panels are sorted by score in descending order (lower scores are better).
Upwards and downwards arrows next to parameter values indicate whether a value is higher or lower than the default parameter.

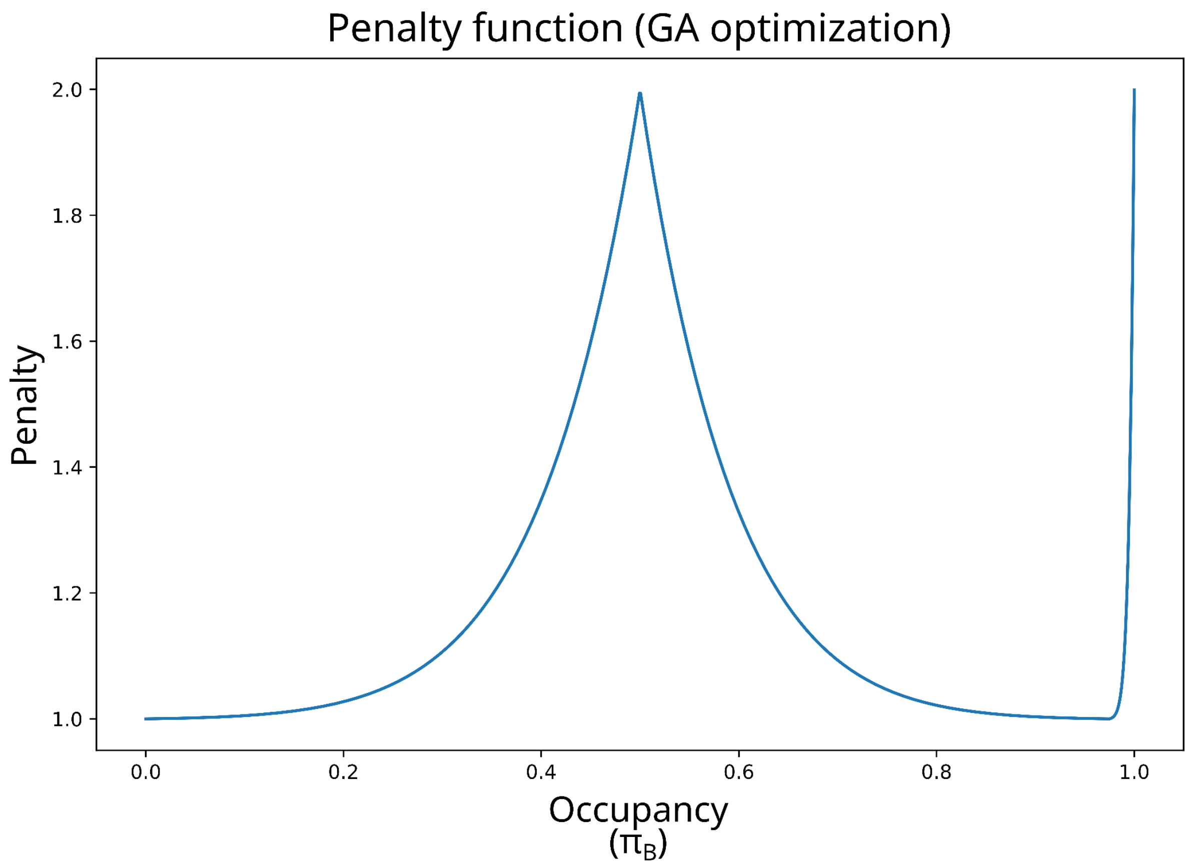


**Supplementary Fig. 12:** Plot of the penalty function used in the GA optimization of extrusion barrier parameters.
