## Additional File 3 for "MoDLE: High-performance stochastic modeling of DNA loop extrusion interactions"

#### Supplementary tables

**Supplementary Table 1**: Parameter table for MoDLE simulations used to generate Fig. 2.

| Figure | Bin size (bp) | Extrusion barrier occupancy | Target contact density | # of LEFs per Mbp | Diagonal width (Mbp) | Avg. LEF processivity (kbp) | Contact sampling strategy |
|---|---|---|---|---|---|---|---|
| 2A | 5000 (default) | Variable | 20.0 | 20 (default) | 3 (default) | 300 (default) | Loop only with noise |
| 2B | 5000 (default) | Variable | 20.0 | 20 (default) | 3 (default) | 300 (default) | TAD only with noise |
| 2C, 2D, 2F (WT) | 5000 (default) | Variable | 20.0 | 20 (default) | 3 (default) | 300 (default) | TAD plus loop with noise (default) |
| 2F ΔCTCF | 5000 (default) | 0.2 | 20.0 | 20 (default) | 3 (default) | 300 (default) | TAD plus loop with noise (default) |
| 2F ΔWAPL | 5000 (default) | Variable | 10.0 | 20 (default) | 6 | 3000 | TAD plus loop with noise (default) |

**Supplementary Table 2**: List of MoDLE software dependencies.

| Name | Project URL | Version | License |
|---|---|---|---|
| Abseil C++ | abseil.io | 20211102 LTS | Apache 2.0 |
| bitflags | github.com/m-peko/bitflags | 1.5.0 | MIT |
| Boost C++ | boost.org | 1.79.0 | BSL 1.0 |
| bzip2 | sourceware.org/bzip | 1.0.8 | BSD-like |

| | | | |
|---|---|---|---|
| | 2 | | |
| Catch2 | github.com/catchorg/Catch2 | 3.1.0 | BSL 1.0 |
| CLI11 | github.com/CLIUtils/CLI11 | 2.2.0 | BSD 3 |
| cmake-git-version-tracking | github.com/andrew-hardin/cmake-git-version-tracking | N/A | MIT |
| concurrentqueue | github.com/cameron314/concurrentqueue | 1.0.3 | Simplified BSD |
| cpp-sort | github.com/Morwenn/cpp-sort | 1.13.0 | MIT |
| fast_float | github.com/fastfloat/fast_float | 3.5.1 | MIT |
| fmt | github.com/fmtlib/fmt | 9.0.0 | MIT |
| HDF5 [1] | hdfgroup.org | 1.12.2 | BSD 3 |
| libarchive | libarchive.org | 3.6.1 | BSD 2 |
| libBigWig | github.com/dpryan79/libBigWig | 0.4.7 | MIT |
| libcuckoo [2,3] | github.com/efficient/libcuckoo | 0.3.1 | Apache 2.0 |
| LZ4 | lz4.org | 1.9.3 | BSD 2 |
| LZO | oberhumer.com/opensource/lzo | 2.1.0 | GPL 2.0 |
| range-v3 | github.com/ericniebler/range-v3 | 0.12.0 | BSL 1.0 |
| readerwriterqueue | github.com/cameron314/readerwriterqueue | 1.0.6 | Simplified BSD |
| spdlog | github.com/gabime/spdlog | 1.9.2 | MIT |
| bshoshany-thread-pool [4] | github.com/bshoshany/thread-pool | 3.3.0 | MIT |
| TOML++ | marzer.github.io/tomlplusplus | 3.1.0 | MIT |
| Xoshiro-cpp | github.com/Reputele | 1.1 | MIT |

| | | | |
|---|---|---|---|
| | ss/Xoshiro-cpp | | |
| xxHash | xxhash.com | 0.8.1 | BSD 2 |
| XZ Utils | tukaani.org/xz | 5.2.5 | GPL 3.0 |
| zlib | zlib.net | 1.2.11 | Zlib |
| zstd | github.com/facebook/zstd | 1.5.2 | BSD 3 |

### References


1. Group HDF, Others. Hierarchical data format version 5. 1997;

2. Fan B, Andersen DG, Kaminsky M. {MemC3}: Compact and Concurrent {MemCache} with Dumber Caching and Smarter Hashing. 10th USENIX Symposium on Networked Systems Design and Implementation (NSDI 13). 2013. p. 371–84.

3. Xiaozhou Li Princeton University, David G. Andersen Carnegie Mellon University, Labs MKI, Michael J. Freedman Princeton University. Algorithmic improvements for fast concurrent Cuckoo hashing [Internet]. ACM Conferences. [cited 2022 Sep 13]. Available from: https://dl.acm.org/doi/10.1145/2592798.2592820

4. Shoshany B. A C++17 Thread Pool for High-Performance Scientific Computing [Internet]. arXiv [cs.DC]. 2021. Available from: http://arxiv.org/abs/2105.00613
